# Non-translated mRNA levels determine P-body properties

**DOI:** 10.64898/2026.03.18.712576

**Authors:** Debdatto Mookherjee, Madeleine Rommel, Ferdinand Weidner, Matej Siketanc, Maria Hondele, Anne Spang

## Abstract

Translational repression enables rapid adaptation to environmental changes. Under stress, translational repressed mRNA and mRNA decay factors accumulate in cytoplasmic processing bodies (PBs), implicated in mRNA storage and decay. PBs have been mostly studied under glucose starvation in yeast, yet, knowledge is limited under other stress conditions. Here, we identify a correlation between the level of translation attenuation and the number, brightness, fluidity and recruitment of PB core components. Stresses triggering strong translation attenuation caused the formation of few bright and more fluid PBs that recruit the decay factors *en bloc*. Conversely, weaker translation attenuation induced numerous, dim, more viscous PBs to which PB proteins were sequentially recruited. Importantly, increasing non-translated mRNA levels augmented the brightness of dim PBs and accelerated decay machinery recruitment. Finally, boosting RNA levels increased the size of Dhh1 helicase-containing droplets *in vitro*. Taken together, we propose a model in which the assembly pathway and biophysical properties of PBs are governed by non-translated mRNA abundance.

**Teaser:** Biophysical properties, protein composition and assembly pathways of processing bodies are dependent on available mRNA levels.

## Introduction

Cells have evolved to adapt to environmental challenges. Coping with stresses requires the cell to rapidly regulate their gene expression on the translational and transcriptional level. Thus, some mRNAs will need to be degraded or stored by the cell. This process is temporally and spatially controlled by the formation of biomolecular condensates like P-bodies (PBs). PBs consist of non-translating mRNAs, the mRNA decay machinery and auxiliary factors, and function as sites of either storage or decay (*1–4*). Stored mRNAs can return to translation thereby maintaining the balance between storage, degradation and translation (*5, 6*).

In *S. cerevisiae*, cytoplasmic mRNA turnover occurs predominantly through the 5′-3′ decay pathway. This process involves sequential deadenylation, decapping, and exonucleolytic degradation from the 5’ end. It is generally assumed that following shortening of the poly(A) tail, the PB components Pat1 and the Lsm1-7 complex bind the shortened poly(A) tail, thereby initiating recruitment and decapping of the mRNA by Dcp1/2 (facilitated by activators such as Edc3 and Dhh1). The uncapped transcript is then subsequently degraded in the 5′ to 3′ direction by the exonuclease Xrn1 (*7–11*).

Beyond decay, the PB components Pat1 and Dhh1 also have direct roles in repression of translation, independent of mRNA decay (*8, 12*). Moreover, inhibition of translation initiation leads to PB formation. Blocking translation elongation by cycloheximide treatment and thereby preventing release of the mRNAs from ribosomes or mutations impede PB formation or lead to their dissolution (*1, 13*). Together, this interplay highlights the integration of translation status and PB dynamics as part of the cellular stress response.

Interestingly, PBs display stress-specific properties. For example, glucose starvation induces a few large PBs per cell, whereas high NaCl and high CaCl_2_ cause the formation of dimmer but more numerous PBs that eventually dissolve due to adaptation (*1, 14*). In addition, defects in the secretory pathway induce multiple dim PBs (*14*). Moreover, it was shown that RNA composition of PBs is context-dependent in yeast and mammalian cells (*15, 16*). These observations raise the question of how PB composition and dynamics are regulated under different stresses. Importantly, the effects of stress specificity on PB properties and translation remains poorly understood.

In this study, we provide evidence that PBs exhibit stress-specific differences in morphology, dynamics, and core PB protein recruitment in *S. cerevisiae* that are linked to the translational state of the cell. Endoplasmic reticulum (ER) stresses (DTT, tunicamycin) weakly inhibit translation, producing dim, viscous PBs with sequential recruitment of core PB proteins. In contrast, conditions that impose stronger translation inhibition (glucose starvation, H_2_O_2_, and CCCP) yield brighter, more fluid PBs to which the decay machinery is recruited *en bloc*. Increasing the pool of non-translating mRNA—via the loss of the polysome-associated proteins Bfr1 and Scp160 or disruption of the deadenylase scaffold Not1—shifted dim PBs towards brighter PBs accompanied by a faster core protein recruitment, directly linking mRNA availability to PB properties. Consistently, elevated mRNA levels enlarged and increased the brightness of Dhh1-containing droplets *in vitro*. Together, our findings support a model in which the abundance of non-translating mRNA dictates PB morphology, assembly pathway, and biophysical properties.

## Results

### Different Stresses Induce PBs with Distinct Number and Brightness

It has been previously reported that PBs differ in number, brightness and half-time depending under which stress they are formed. While glucose starvation-induced PBs are bright, large and persist until glucose replenishment, PBs induced by hyperosmotic stresses form multiple and small PBs that dissolve within 30-40 min of stress (*14*). To investigate the general principles underlying these differences, we employed five different physiologically relevant and PB-inducing stress conditions at acute (10 min) and prolonged (60 min) exposure times. The selected stresses cover metabolic stress (glucose starvation), redox stress (DTT and H_2_O_2_), ER stress (DTT and tunicamycin) and mitochondrial dysfunction (CCCP) (Figure 1A). The concentration of the chemical stressors was chosen to not induce cell death (Figure S1A). Using the decapping enzyme Dcp2-GFP as a PB marker, we observed differences in number and brightness of PBs depending on the stressor (Figure 1B). As previously reported, glucose starvation induced ∼2 bright PBs per cell. H_2_O_2_- and CCCP-induced PBs were moderately dimmer and more numerous, while DTT- and tunicamycin-induced numerous rather dim PBs (Figure 1C-G). The number of PBs per cell upon CCCP, DTT and tunicamycin treatment was comparable. Thus, there appears to be no strict correlation between brightness and number of PBs upon stress induction. Notably, the stressors DTT and Tunicamycin caused a slower onset of PB formation as in about 20% of cells PBs were not detectable within 10 min of stress (Figure S1B-C). Under glucose starvation, PB and SG formation was shown to be coupled (*6, 17, 18*). Therefore, we checked for the formation of SGs under our stress conditions at acute and prolonged time points using two different markers Pub1-GFP and Tif4632-GFP. However, we could not observe the formation of SGs (Figure S1D), consistent with the notion that SG induction and PB induction is mostly uncoupled in *S. cerevisiae* (*19*). It is conceivable that the increased brightness of PB correlates with the strength of the stressor. This, however, does not appear to be the case. On the contrary, both, the very dim PB-inducing stress DTT and glucose starvation, which induces very bright PBs, lead to the strongest growth attenuation (Figure 1H). Hence, there is no correlation between brightness, number and strength of the stressor in PB formation.

**Figure 1.**
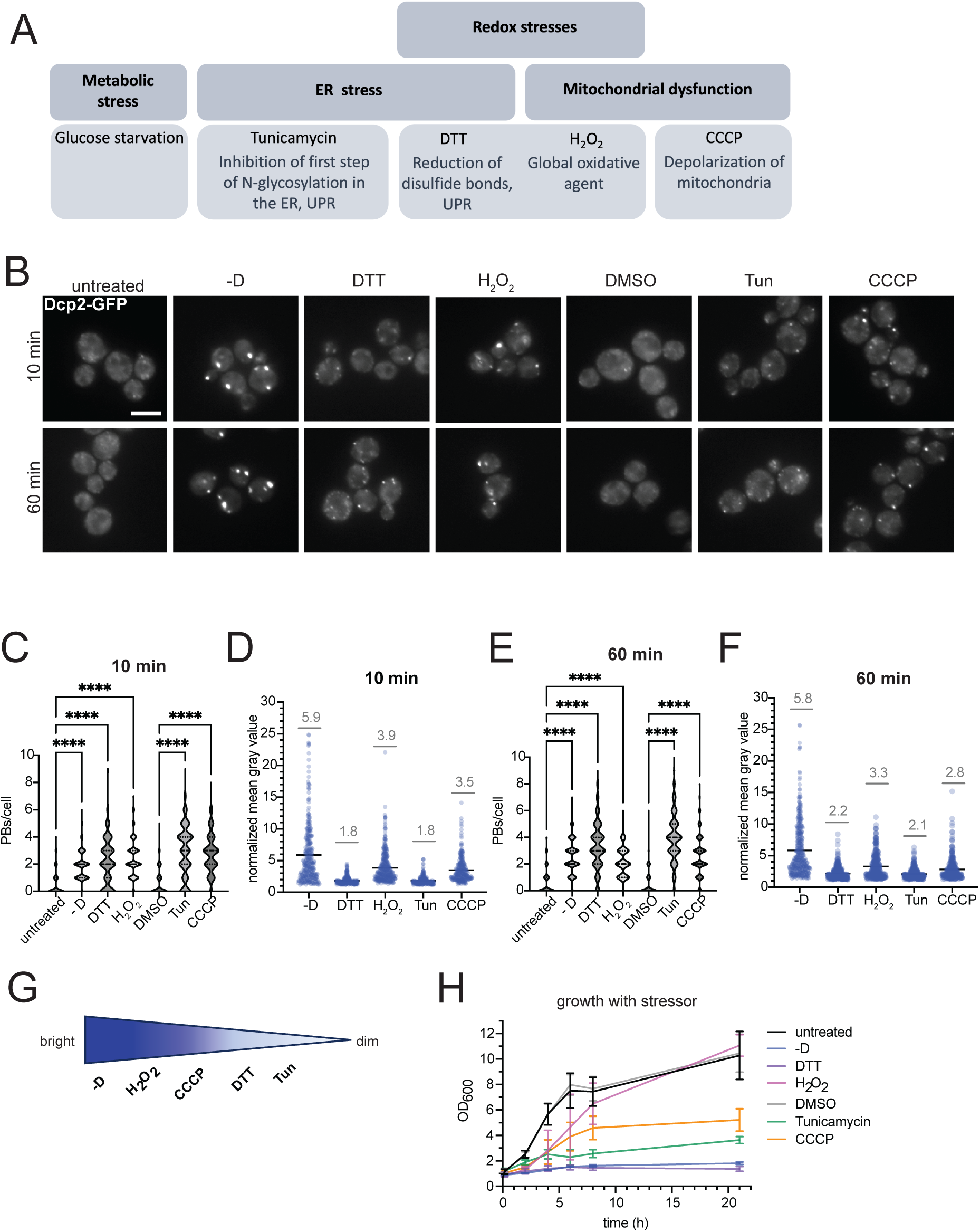
Different stresses induce PBs with different brightness and number. (A) Overview of the repertoire of stresses that were used to induce PBs. (B) Cells expressing chromosomally GFP-tagged Dcp2 were starved for glucose or treated with 50 mM DTT, 500 µM H_2_O_2_, 10 µg/ml tunicamycin and 20 µM CCCP for 10 min or 60 min. DMSO is the vehicle control for CCCP and tunicamycin. n=3. Scale bar = 5 µm (C) Depicts the number of PBs per cell after 10 min of stress. The fluorescence intensity of the PBs is plotted in (D). (E) Shows the PBs per cell after prolonged stress (60 min) and the (F) fluorescence intensity of PBs after 60 min of stress. (G) Represents the brightness index of the PBs under different stresses. (H) Yeast cells were treated with the stressor and their growth was observed by tracking OD_600_ for 21h in n=3 independent experiments. Statistical analysis was performed using a Kruskal-Wallis test with Dunn’s multiple-comparisons test in GraphPad Prism. \*\*\*\**p* < 0.0001.

### Biophysical properties of PBs are stress-dependent

Given the differences in morphology and number of the PB, we were next exploring how they would dissolve after stress relief. As expected, we observed immediate dissolution of PBs after glucose replenishment. A similar effect, albeit with a slight delay, we also noticed after washout of H_2_O_2_ and CCCP. In contrast, PBs induced by DTT and tunicamycin persisted for at least 2 h after stress washout (Figure 2A-B). To exclude the possibility that removal of DTT or tunicamycin was not successful and this would cause the persistence of the PBs, we first recorded the growth after stress removal. After DTT washout, cells resumed growth with similar kinetics than for the other stressors. After tunicamycin washout, cells only slowly started to grow again, suggesting that the stress persisted in those cells (Figure 2C). Both DTT and tunicamycin elicit the unfolded protein response (UPR). Therefore, we determined next the UPR state under those stresses by detecting the level of spliced HAC1 mRNA (Figure S2A). Consistent with the growth restoration, UPR was turned off about 30 min after DTT washout. After tunicamycin washout, however, cells retained spliced HAC1 throughout the time course, indicating that UPR was still on and this might be the reason for the slowed growth rate after stress removal (Figure 2D). Taken together, our data suggest the presence of temporary (-D, H_2_O_2_, CCCP) as well as persistent PBs (DTT) and that the brightness of PBs might be predictive of the biophysical properties (Figure 1G and Figure 2E).

**Figure 2.**
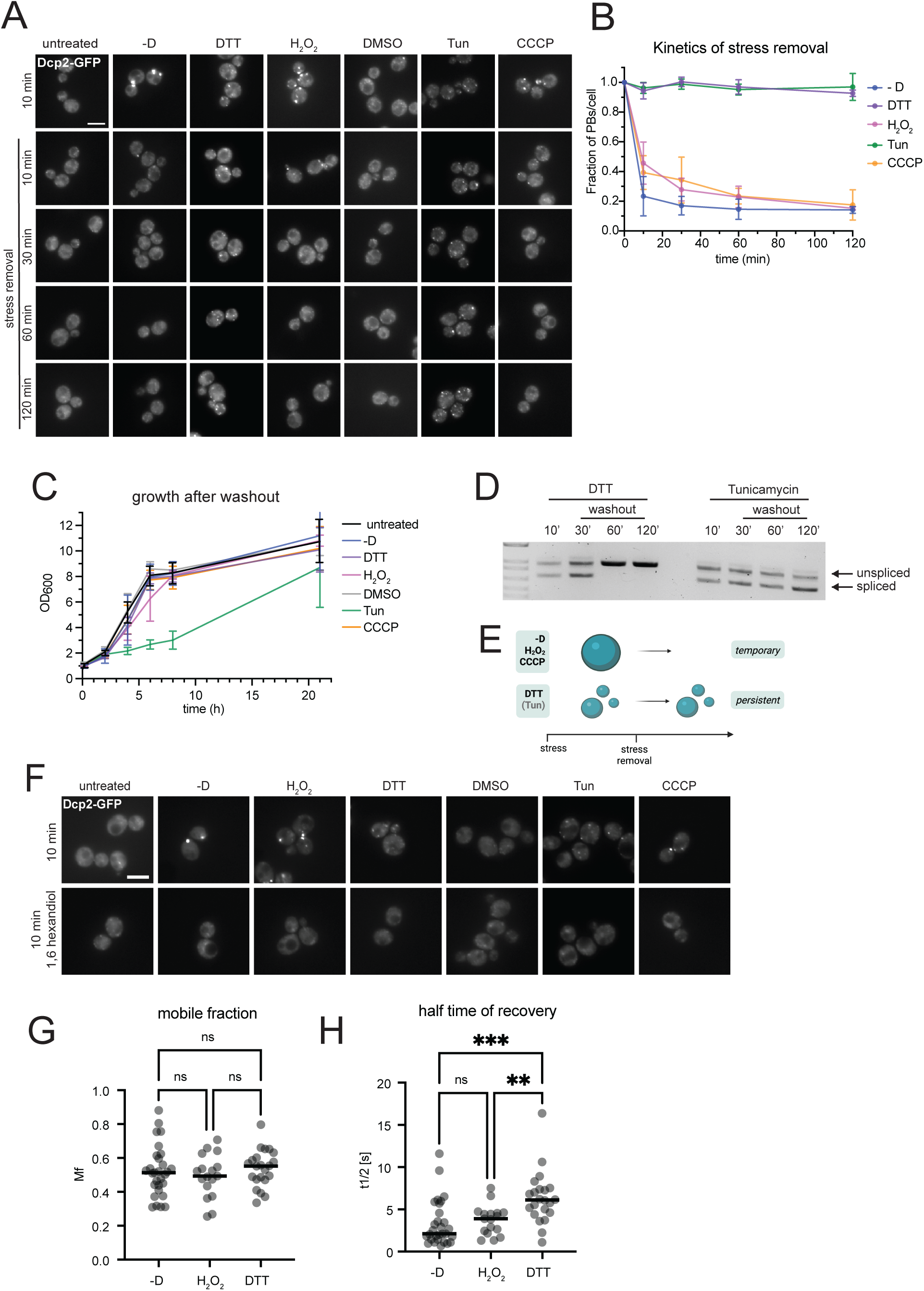
DTT and Tunicamycin induce persistent PBs. (A) Yeast cultures were treated with the stressors for 10 min, spun down, washed and resuspended in YPD. PBs (Dcp2-GFP) were tracked for 10, 30, 60 and 120 min after stress removal. n=3. Scale bar = 5 µM (B) The PBs per cell were quantified using thresholding algorithms of ImageJ and were normalized to the number of PBs at 10 min of stress. PBs induced by tunicamycin and DTT do not wash out after stress removal. (C) Ensuring a successful drug washout, growth after stress removal was determined by tracking OD_600_ for 21h. n=3. (D) UPR activation and deactivation levels were established by a HAC1 splicing assay. Stress was induced for 10 min, and samples were taken at timepoints after stress removal 30, 60 and 120 min. n=3. (E) Schematic representation of stresses (-D, H_2_O_2_ and CCCP) that form temporary PBs and stresses (DTT and tunicamycin) that form persistent PBs. (F) 1,6-hexandiol dissolves PBs induced under a variety of stresses. Yeast cells were treated with or without 1M 1,6-hexandiol for 10 min after stress induction for 10 min. n=3. Scale bar = 5 µm (G) FRAP analysis of PBs induced under different stresses. Yeast cells expressing chromosomally GFP-tagged Dhh1 (Dhh1-GFP) were stressed for 30-60 min and then photo-bleached. Their fluorescence recovery was measured over 10 s and the mobile fraction (Mf) was determined using FrapBot. The experiment was repeated for -D and H_2_O_2_ in n=3 and for DTT in n=4 individual experiments. One-way ANOVA with Tukey’s multiple-comparisons test was used for statistical analysis. (H) Depicts the half time of recovery (t_1/2_) of at least 19 PBs per condition that were calculated using FrapBot webtool. Kruskal–Wallis test with Dunn’s multiple comparisons test was used for statistical analysis. ns, not significant, **p < 0.01, ***p < 0.001.

However, at this point we were not able to exclude that the persistent DTT-induced PBs represent aggregates, which are not dissolvable. To address this question, we first treated cells with 1,6-hexandiol, which is known to dissolve membraneless organelles (*20*). Indeed, temporary as well as persistent PBs dissolved within 10 min of 1,6-hexandiol treatment, indicating that the persistent PBs were not aggregates (Figure 2F). Next, we used FRAP to detect differences in their liquid-like properties in a strain with chromosomally GFP-tagged Dhh1. We could not detect any significant difference in the mobile fraction between temporary (-D and H_2_O_2_) and persistent (DTT) PBs, confirming that persistent PBs were not mere aggregates. However, DTT-induced PBs showed slightly slower diffusional exchange, implying a moderately higher viscosity compared to temporary PBs (Figure 2G-H). Taken together, PBs formed under different stresses can vary in their biophysical properties, which potentially may contribute to their lifetime after stress removal.

### Persistent PBs Show a Delay in Recruitment of 3’ Protein Components

We have shown above that the DTT-induced PBs have different properties compared to temporary PBs. This difference in properties could be driven by the protein composition, the mRNA content, or both. To distinguish between these possibilities, we first determined the recruitment of core PB protein components over time under various stresses. To this end, we created double-tagged strains, expressing endogenously Dcp2-GFP and individually other PB components tagged with mCherry, namely Dcp1, Edc3, Dhh1, Xrn1, Pat1 and Lsm4. As had been previously reported for glucose starvation, core PB protein components were recruited *en bloc* (*21*) (Figure 3A-B). We observed a similar *en bloc* recruitment for the two other stressors causing temporary PBs, H_2_O_2_ and CCCP. In contrast, we found that the persistent PBs were unable to recruit 3’ components within 10 min of stress, but rather display a staggered recruitment of PB core components, starting from the 5’ end towards the 3’ end (Figure 3C-D, Figure S3A-B). We quantified the protein recruitment manually at selected timepoints. To get a higher temporal resolution and to independently confirm our observations, we employed an unbiased, automated imaging approach combined with AI-assisted PB detection and determined the sequence of recruitment. This approach confirmed our findings (Figure 3E-F and Figure S4A-H). In both analyses, recruitment of the 3′ components were delayed under stresses that induced persistent PBs. While the manual and AI methods differed somewhat in the absolute timing of recruitment, both quantification approaches showed the same consistent trends, supporting delayed recruitment from the 5’ to the 3’ as a reproducible feature of PBs induced by persistent stress. Taken together, our data indicate that persistent PBs assemble more slowly than temporary PBs, with a staggered and delayed recruitment of 3’ core components (Figure 3G), and suggest that distinct assembly dynamics may impact their functional capacity and also at least in part the difference in biophysical properties. Thus, the difference in protein composition of PBs under various stresses may contribute to the different biophysical properties.

**Figure 3.**
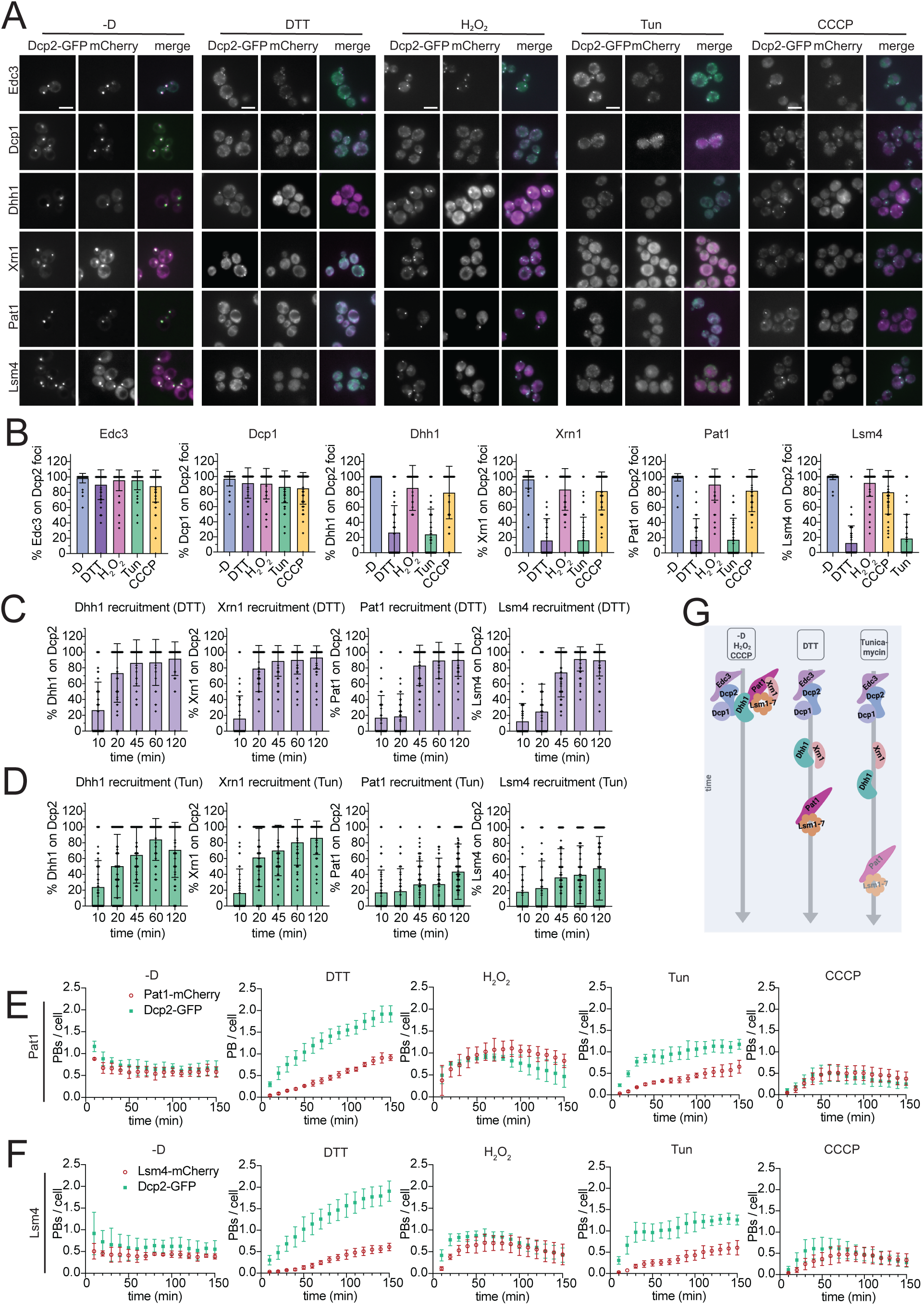
Core PB proteins are recruited sequentially to persistent PBs. (A) Dcp2-GFP tagged cells were also chromosomally mCherry tagged with Dcp1, Dhh1 Xrn1, Pat1 or Lsm4. PB formation was assessed after 10 min of stress. n=3. Scale bars = 5 µM. (B) Quantification of the the percentage of mCherry tagged proteins overlapping with the Dcp2-GFP signal per cell per condition after 10 min of stress. Mean and standard deviation are shown for 90 cells from n=3 independent biological replicates. (C) 3’ PB components are recruited over time under DTT stress. Colocalization was quantified as the percentage of mCherry-tagged proteins overlapping with Dcp2-GFP per cell after different times of DTT stress. Data are presented as mean ± standard deviation from 90 cells from n=3 independent biological replicates. (D) 3’ PB components are recruited over time under tunicamycin stress. Quantification of the percentage of mCherry-tagged proteins colocalizing with Dcp2-GFP per cell after different times of tunicamycin treatment. Values represent the mean ± standard deviation of 90 cells from n=3 independent biological replicates. (E) AI-assisted time-resolved quantification of PB per cell in Dcp2-GFP/Pat1-mCherry or (F) Dcp2-GFP/Lsm4-mCherry expressing cells. Cells were imaged automatically in 384-well plates, and PB were quantified using Nikon NIS AI modules. Data represent mean ± standard deviation from n=3 independent biological replicates, with ≥250 cells quantified per timepoint and replicate. (G) Schematic representation of *en bloc* versus sequential recruitment to PBs of the different stress.

### Levels of Translation Attenuation Correlate with PB Brightness

Even though we showed above that the core PB protein recruitment pathway under various stresses may contribute to the different PB properties, this does not exclude a contribution by the mRNA. Therefore, we addressed next the role of mRNA in PB assembly. Translation and PB formation are tightly linked. To test whether stress-specific properties such as brightness, viscosity and PB core protein recruitment, are linked to the amounts of non-translated mRNA available for recruitment into PBs, we determined the level of translation under different stresses using an L-HPG incorporation assay. The methionine analog L-HPG is integrated into newly synthesized proteins and can be conjugated to the fluorophore picolyl-azide AF488 via click chemistry. This fluorescence serves as a direct readout of translation levels and is measured by FACS (Figure 4A). As expected, 10 min of glucose starvation led to severe translation attenuation (*12, 14, 22*). A similar effect was observed for H_2_O_2_ treatment (Figure 4B-C). CCCP caused a major translational drop of 60-70%, followed by DTT with about 30%. Surprisingly, cells treated with tunicamycin did not show significant translation attenuation. Therefore, we repeated the experiment after 60 min stress exposure. Unexpectedly, even prolonged tunicamycin treatment did result only in small but insignificant translation attenuation, while prolonged exposure to DTT further decreased the translational capacity (Figure 4B-C). Therefore, we thought of an independent manner to assess translational attenuation and determined the levels of phosphorylated eIF2*α* (Figure S5A-B). The western blots confirmed our L-HPG incorporation results in that the strongest translation attenuation occurred under H_2_O_2_ treatment, followed by CCCP and DTT. Again, we failed to detect translation attenuation when cells were treated with tunicamycin. Of note, although the translation attenuation under glucose starvation is very strong, the response is not regulated via eIF2*α phosphorylation, but leads to the dissociation of eIF4 from translation initiation complex (22).* We were puzzled by the apparent lack of translation attenuation upon tunicamycin treatment. It is conceivable that our methods are not sensitive enough to detect minor changes in translational capacity. To test this possibility, we treated cells with cycloheximide (CHX), which locks ribosomes on mRNAs and is a potent inhibitor of PB formation under glucose starvation (*1, 23, 24*). Application of CHX before the stressors, inhibited PB formation under all stress conditions, demonstrating that mRNA release from polysomes is necessary for PB formation also upon tunicamycin treatment, even though we observed no significant translation attenuation in tunicamycin-treated cells using the L-HPG assay or eIF2*α* phosphorylation levels (Figure 4D). Thus, minor mRNA release from the translating pool is sufficient to initiate PB formation. In addition, we added CHX to cells after PB formation under the different stresses and observed dissolution of all temporary and persistent PBs. These results suggest that PBs need constant influx of mRNA from translational pool for their maintenance (Figure 4E-F). Our data also demonstrate a correlation between the brightness of the PBs and the level of translation attenuation and thereby the level of non-translated mRNA in the cytoplasm. Thus, we hypothesized that the level of non-translated mRNA available for sequestration into PBs might at least in part determine the different PB properties such as brightness and the assembly pathway.

**Figure 4.**
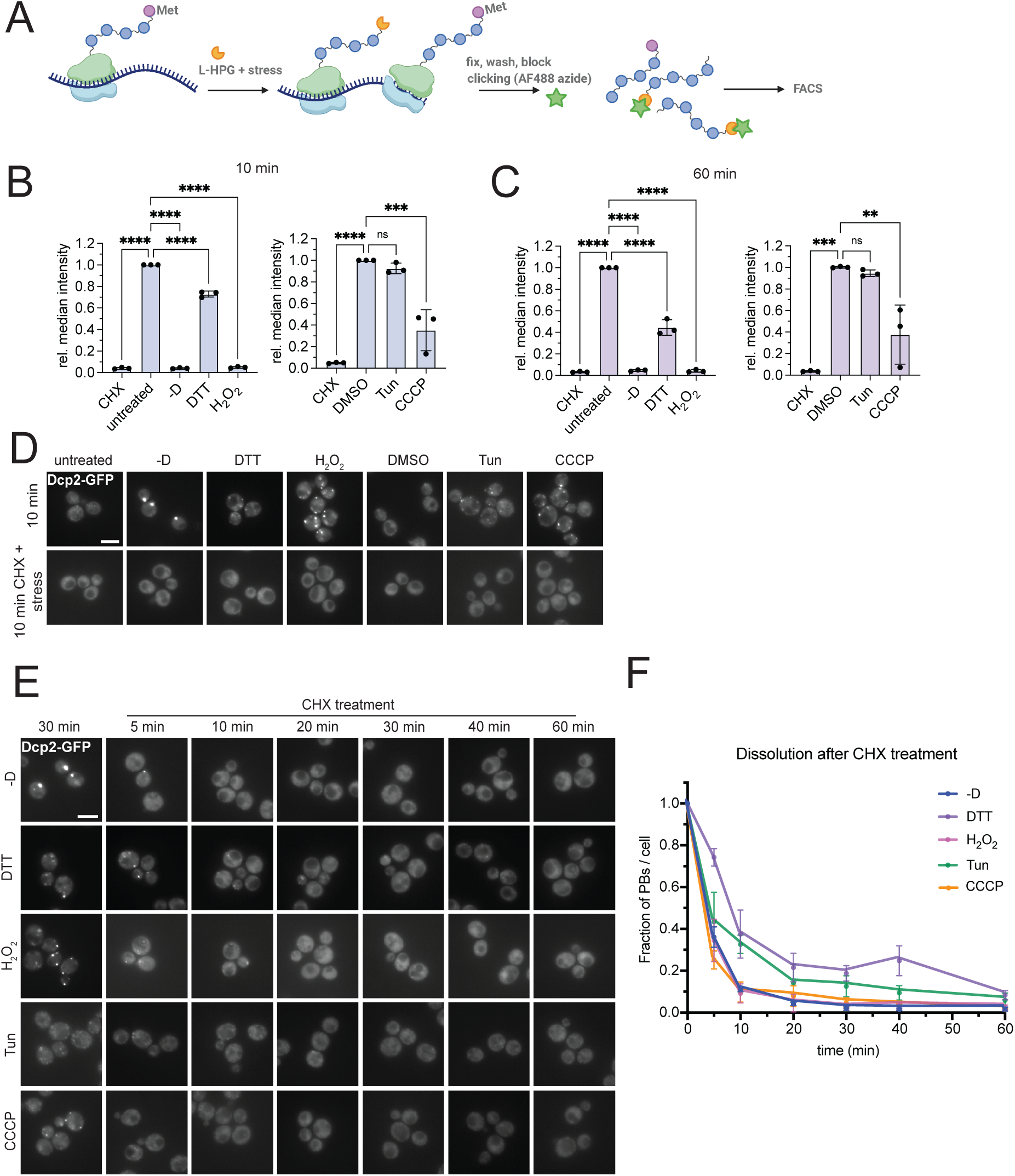
Translational state of the cell correlates with PB brightness. (A) Schematic representation of the L-HPG translation assay, where L-HPG is added to a methionine free medium for the time of the stress. L-HPG, which contains an alkyne moiety, is incorporated into the nascent peptide. After fixation, washing and blocking of the fluorescent dye AF488-picolyl azide can be clicked to the incorporated L-HPG. Fluorescence intensities are measured by FACS. (B) Quantification of global translation as measured by using L-HPG. Different stress conditions normalized to the median fluorescence intensity of the vehicle control (untreated or DMSO) after 10 min or (C) 60 min of stress were compared for n=3 independent biological experiments. One-way ANOVA with Tukey’s multiple-comparisons test was used for statistical analysis. ns, not significant*, \*\**p *< 0.01*, *\*\*\**p *< 0.001, \*\*\*\**p *< 0.001.* (D) Widefield microscopy of cells that were stressed for 10 min with or without CHX pretreatment for 10 min. n=3. Scale bar = 5 µm. (E) Cells were stressed for 10 min before the addition of 100 µg/ml CHX. Images were taken 5, 10, 20, 30, 40 and 60 min after CHX addition. For comparison, cells were imaged after 30 min addition of the stressors. n=3. Scale bar = 5 µm. (F) Quantification of the change of PB number per cell over time normalized to the number of PBs after 30 min of stress.

### The Polysome-associated Proteins Bfr1 and Scp160 Regulate Properties of DTT PBs

To test our hypothesis, we sought ways to genetically increase the levels of non-translated mRNAs in the cytoplasm. We have shown previously that under normal growth conditions, deletion of polysome-associated proteins, namely Bfr1 and Scp160 negatively regulate PB formation (*25*). Deletion of Scp160 led to the formation of pseudo-PBs, containing Dcp2 and partially Edc3 and Pat1, but not Dhh1, while *bona fide* PBs were formed in the absence of Bfr1. Thus, Bfr1 and Scp160 were suggested to protect mRNAs from accessing PB components (*25*). Hence, deletion of the *BFR1* and *SCP160* would increase the amount of non-translated mRNAs in the cytoplasm. In support of this notion, it was previously shown that Scp160 is necessary for the efficient translation of codon-optimized mRNAs. In the absence of Scp160, these transcripts may not be effectively engaged by ribosomes, leading to slower translation, reduced ribosome association, consequently rerouting into RNA granules such as PBs (*26*). DTT-induced PBs were brighter and more numerous in *Δbfr1 Δscp160* cells compared to wild type (Figure 5A-C). In addition, those DTT-induced PBs in *Δbfr1 Δscp160* dissolved rapidly after DTT washout indicating a change in biophysical properties of the PBs (Figure 5D-E). Moreover, in *Δbfr1 Δscp160* cells, the recruitment of the PB components Pat1 and Lsm4 was accelerated, indicating a faster PB assembly pathway (Figure 5F-I). These results support our hypothesis that an increase of non-translated mRNA changes the PB properties. However, deletion of either *SCP160* or *BFR1* cause polyploidy. Therefore, we repeated the experiment and only depleted Scp160 using an auxin inducible degron. Scp160 was successfully depleted after 120 min of 100 µM indole-3-acetic acid (IAA) treatment (Figure S6 A-B). Depletion of Scp160 for 2 h did not lead to polyploidy (*26, 27*). The DTT-induced PBs in Scp160-depleted cells still showed increased brightness, albeit to a lesser extent than in the *Δbfr1 Δscp160* double mutant (Figure S6C-D). Moreover, those PBs were at least partially dissolvable after stress removal (Figure S6E-F). Our data so far support our hypothesis that the amount of mRNA released from the translational pool dictates the brightness, assembly pathway and biophysical properties of PBs.

**Figure 5.**
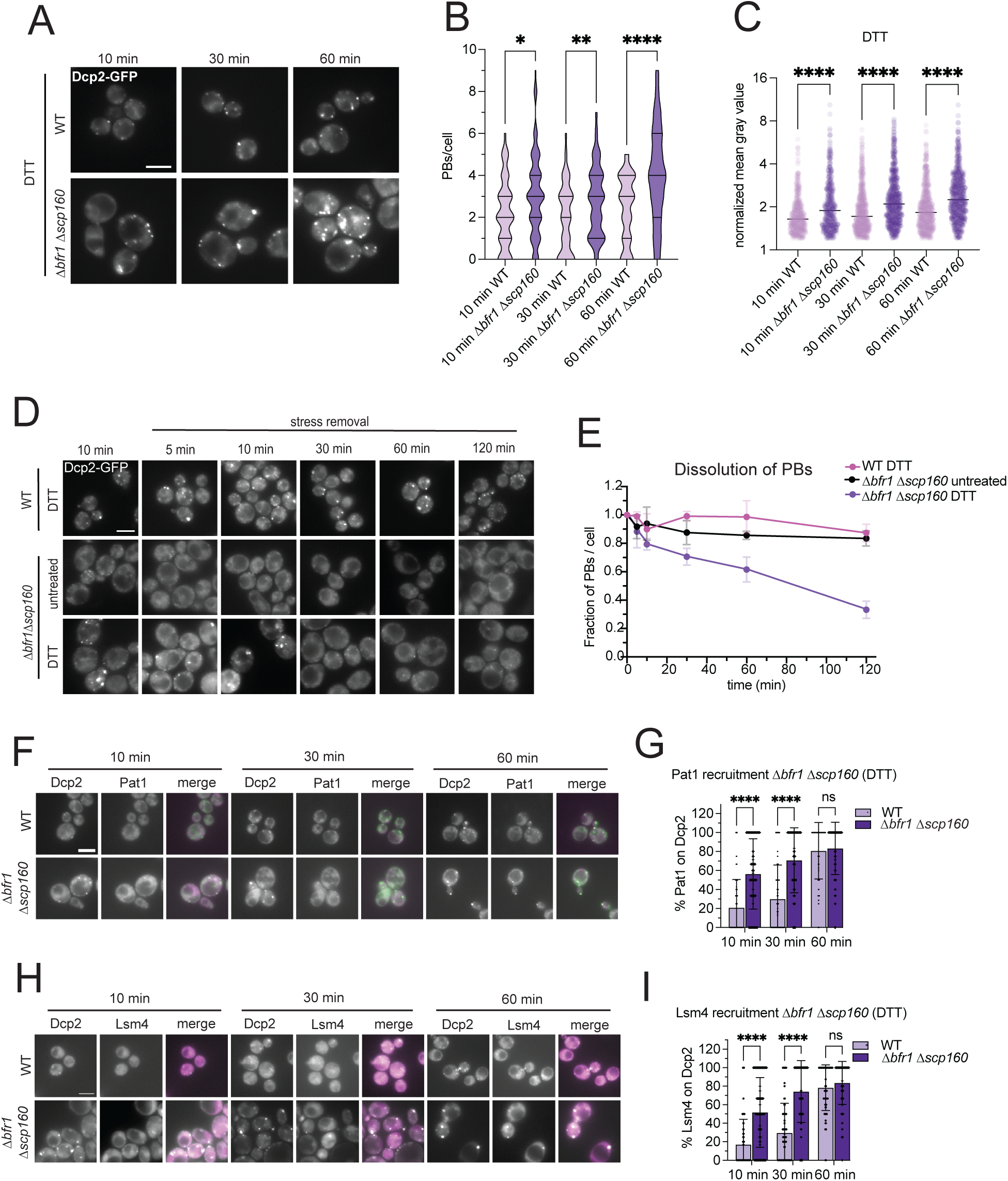
Deletion of the polysome-associated proteins Bfr1 and Scp160 causes changes of DTT PB properties. (A) Wild-type and *Δbfr1Δscp160* cells expressing Dcp2-GFP were imaged 10, 30 and 60 min after DTT addition. n=3 Scale bar = 5 µm. (B) Quantification of PBs per cell in WT and *Δbfr1Δscp160* cells. (C) Analysis of the fluorescence intensity of PBs after 10, 30 and min of stress in wild-type and *Δbfr1 Δscp160* cells. The y-axis is displayed on a log_2_ scale. Kruskal–Wallis test followed by Dunn’s multiple comparisons test was used for statistical analysis of (B) and (C). *p < 0.05, **p < 0.01, ****p < 0.0001. (D) DTT-induced PBs were imaged up to 120 min after washout of the stressor. Untreated *Δbfr 1Δscp160* cells served as a control. n=3. Scale bar = 5 µm. (E) Quantification of PB number per cell shown as depicted change relative to 10 min of DTT treatment. (F) Recruitment of Pat1-mCherry, and (H) Lsm4-mCherry to Dcp2-GFP foci after 10, 30 and 60 min of DTT treatment in WT and *Δbfr1 Δscp160* cells. n=3. (G, I) Quantification of the data shown in (F) and (H). Data were analyzed using two-way ANOVA followed by multiple-comparisons testing with Šídák’s correction. Scale bar = 5 µm. ns, not significant, *\*\*\*\*p < 0.0001*.

### Non-Translated mRNA Levels impact PB Assembly Pathways and PBs’ Biophysical Properties

To corroborate our findings above, we sought a second way to enhance the levels of non-translated mRNA in the cytoplasm in a more acute manner. For that reason, we affected deadenylation, a pre-requisite for mRNA decay. The Ccr4-Not complex is a central regulator of mRNA deadenylation and translation-decay coupling, and perturbation of Not1 is known to increase cytoplasmic non-translated mRNA pools (*28*)(*29–34*). Not1 is essential and therefore, we depleted Not1 using an auxin-inducible degron. Not1 is efficiently depleted after 60-120 min of auxin treatment (Figure S6G-H). Upon Not1 depletion, we observed PB formation, consistent with the notion that PBs can act as mRNA sequestering compartments (Figure 6A-B). Combining DTT stress and Not1-depletion increased the brightness of the PBs compared to non-depleted cells, indicating that the amount of mRNA determines the brightness of the PBs (Figure 6C-E). Moreover, the recruitment of the 3’ components, Pat1 and Lsm4, to DTT PBs was accelerated upon Not1-depletion (Figure 6F-I). We used two types of *in vivo* manipulations, one acute and one permanent, to show that the level of non-translated mRNA determines the brightness and the kinetics of the assembly pathway.

**Figure 6.**
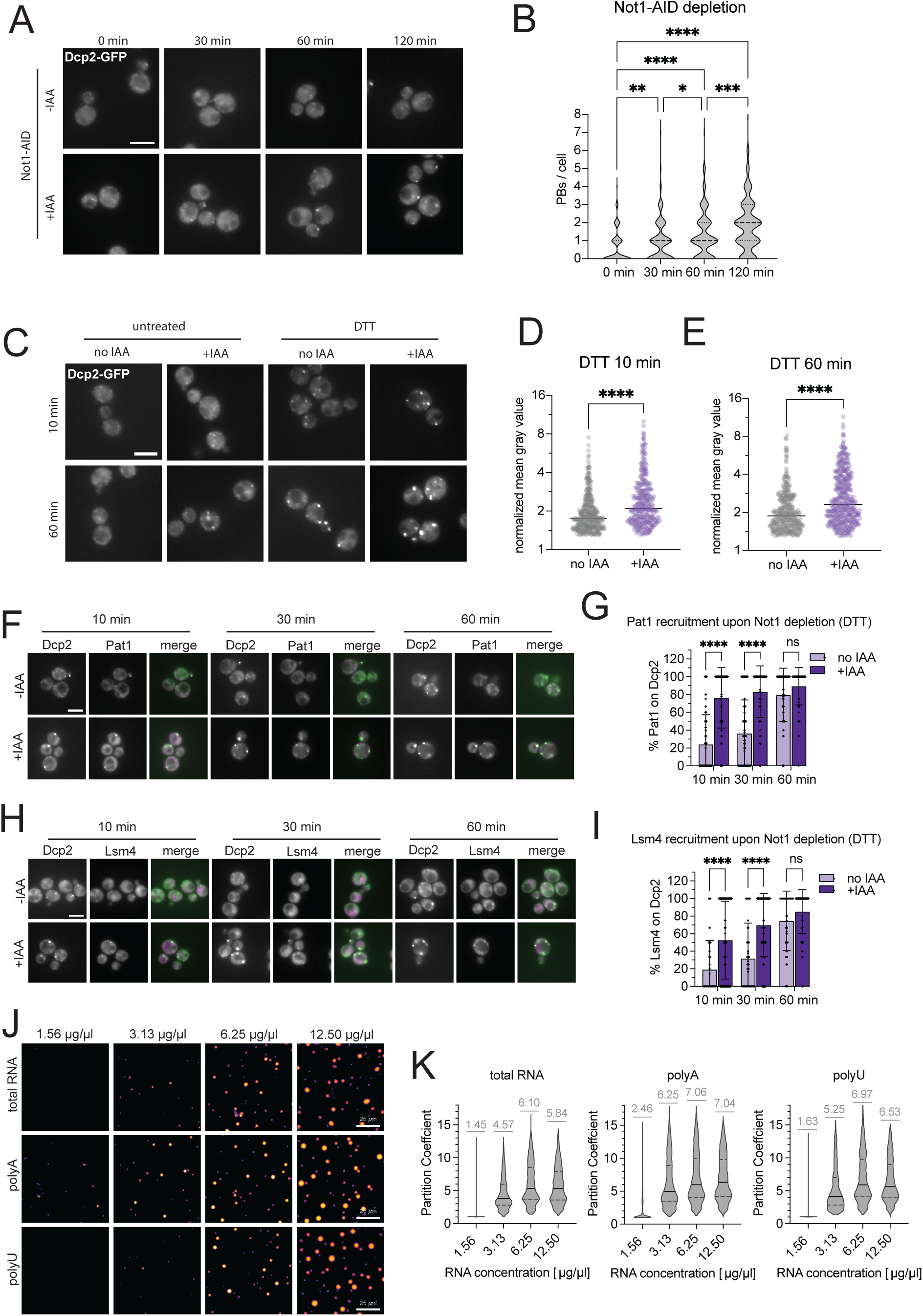
Non-translated mRNA determine PB properties in cells and in vitro. (A) Imaging of PB formation (Dcp2-GFP) upon Not1-depletion up to 120 min. n=3. Scale bar = 5 µM. (B) Violin plots showing the number of PBs per cell, which were quantified using thresholding algorithms of ImageJ. Statistical analysis was performed using a Kruskal–Wallis test with Dunn’s post hoc multiple-comparisons test. *p < 0.05, **p < 0.01, ***p < 0.001, ****p < 0.0001. (C) Imaging of PBs (Dcp2-GFP) in untreated and DTT-treated cells (10 and 60 min), comparing Not1-AID depleted cells (100 µM IAA) to non-depleted cells. Not1 was depleted for 60 min previous to stress treatment and the IAA was kept during the stress treatment. n=3. Scale bar = 5 µm. Quantification of the brightness of PBs after (D) 10 min and (E) 60 min DTT treatment in Not1-AID depleted (+IAA) and non-depleted (no IAA) cells. The y-axis is displayed on a log₂ scale. Statistical significance between the two groups was assessed using a **two-tailed Welch’s t-test.** ****p < 0.0001. (F) Recruitment over time of Pat1-mCherry to Dcp2-GFP foci in Not1-depleted (+IAA) and non-depleted cells (-IAA). Scale bar = 5 µm. (G) Quantification of colocalization Pat1-mCherry foci with Dcp2-GFP foci in Not1-depleted and non-depleted at indicated timepoints. Data were analyzed using two-way ANOVA followed by multiple-comparisons testing with Šídák’s correction. n=3. (H) Recruitment of Lsm4-mCherry to PBs marked by Dcp2-GFP in Not1-depleted (+IAA) and control (-IAA) cells at indicated time points. n=3. Scale bar = 5 µm. (I) Quantification of colocalization Lsm4-mCherry foci with Dcp2-GFP foci in Not1-depleted and non-depleted at the indicated timepoint. Data were analyzed using two-way ANOVA followed by multiple-comparisons testing with Šídák’s correction. n=3. ns, not significant, ****p < 0.0001. (J) *In vitro* phase separation assay of Dhh1 (spiked with 1% Atto565 labelled protein) and titration of increasing amounts of yeast total RNA, polyU and polyA RNA in n=3 independent experiments. (K) Partition coefficient of Dhh1 upon titration of indicated RNA types and concentration.

If the critical determinant in PB properties is indeed the level of available mRNA, we should be able to reconstitute this effect in a minimal *in vitro* system. We therefore used the helicase Dhh1, which has been shown previously to form LLPS in the presence of mRNA (*13, 35, 36*). Keeping a constant concentration of Dhh1, we increased the level of RNA. The brightness and size of the Dhh1 droplets correlated with the increase of RNA in the assay (Figure 6J-K). Together, these findings demonstrate that mRNA availability is a critical determinant of PB formation and their properties, and that RNA itself is sufficient to modulate PB properties *in vivo* and *in vitro*.

## Discussion

Despite observations that PB morphology and number can vary dependent on the stressor, the underlying mechanism remained unresolved (*1, 14, 15*). Here, we systematically compared metabolic, redox, ER, and mitochondrial stresses. We demonstrate that PB number, brightness, and assembly kinetics are stress-specific and do not correlate with overall stress severity as measured by growth inhibition. Glucose starvation and DTT both strongly impair growth, yet induce PBs with diametrically opposed properties—few, bright PBs versus numerous, dim ones, respectively. In addition, we found that the biophysical properties as well as the assembly pathways were different under these stressors. Finally, translation attenuation levels were strikingly different between glucose starvation and DTT treatment. These observations argue against a simple stress-strength model as predominant determinant of PB properties.

We further show that PB life-time after stress removal differs markedly between stresses. PBs induced by glucose starvation, oxidative stress, or mitochondrial dysfunction rapidly dissolve, whereas ER stress-induced PBs persist for hours, even after UPR deactivation in the case of DTT. FRAP and 1,6-hexanediol sensitivity demonstrate that all PBs remain liquid-liquid phase-separated entities. Persistent PBs, however, exhibit slower molecular exchange, consistent with increased viscosity. It has been shown previously that PB properties are modulated by ATP-dependent remodeling and RNA-protein interactions (*13*). Consistent with this observation, our data suggest, that the concentration of mRNA in PBs contributes to the fluidity of the PBs. Based on our results, we propose a model in which cytoplasmic non-translated mRNA concentration and flux define PB assembly pathway, composition, and physical state (Figure S7).

It had been previously assumed that PBs form rapidly under stress by the coalescence of mRNAs on which the decay machinery and the core PB components are already assembled (*1, 2, 37, 38*). While this assembly pathway is likely to act on glucose starvation, H_2_O_2_ and CCCP treatment, our study revealed a second assembly pathway in which the core PB protein components are sequentially recruited from the 5’ to the 3’ end. Although incomplete core PB protein recruitment has been observed previously (*21, 25*), the underlying reason was not addressed. We provide strong evidence for a link between the stress-dependent translational attenuation and core PB protein recruitment. Given that Pat1-Lsm1-7 and Xrn1 are critical for productive 5′-3′ decay (*10, 21*), mRNA decay may be postponed in those PB until the assembly is complete. Therefore, PBs formed under acute DTT or tunicamycin stress are likely to serve as storage compartments rather than promoting mRNA decay. Once the 3’ protein components are recruited, it is conceivable that the PBs at that point are also involved in mRNA decay. It will be interesting to see in the future, whether mRNAs stored in these PBs after acute stress are protected from decay. We have shown previously that even under glucose starvation a subset of mRNAs is protected in PBs from decay (*15*).

Consistent with this RNA-availability model, increasing the pool of non-translated mRNA—either by removal of the polysome-associated proteins Bfr1/Scp160 or by depletion of the Ccr4–Not complex scaffold Not1—shifts ER stress-induced PBs toward a brighter phenotype with accelerated recruitment of core PB components. These genetic perturbations therefore connect translational control and mRNA turnover capacity to PB physical state and assembly kinetics. Mechanistically, an important implication is that PB core protein recruitment should be viewed as modular rather than obligatorily fixed. This possibility receives support from mounting evidence for multiple compositionally distinct decapping assemblies and regulated decapping-factor engagement in yeast (*10, 11*).

Finally, our *in vitro* reconstitution shows that elevating RNA concentration alone is sufficient to increases the size and brightness of Dhh1-containing droplets, reinforcing the view that RNA could also act as a structural driver of PB-like condensates rather than only cargo (*1*). Our findings are consistent with emerging principles of RNA-mediated phase separation, in which RNA concentration, length, and sequence composition and/or structure modulate condensate composition and material state (*39–42*).

Together, our results support a cohesive framework in which stress-specific translation states and as a consequence non-translated mRNA levels tune PB assembly mode and biophysical properties, providing a unifying model for divergent PB phenotypes across conditions.

## Material and Methods

### Yeast methods

Standard genetic techniques were employed throughout the study (*43*). Unless indicated otherwise all genetic mutations were introduced chromosomally. Chromosomal tagging and deletions were carried out as described in (*44–46*). For C-terminal GFP tagging we use pYM-yeGFP-KlTRP1, for mCherry tagging we used pFA6a-3mCherry (*hphNT1*) plasmid and for tagging with auxin-inducible degron we used pHyg-AID*-6HA and pNAT-AID-9myc. Deletions of *BFR1* were introduced using pUG73 (LEU). Strains used are listed in Table S1.

### Yeast cultures and drug treatments

Cells were grown in YPD medium to OD_600_ 0.6-0.8 at 30°C and exposed to the respective stress conditions and controls, 0% glucose, 50 mM DTT, 500 µM H_2_O_2_, 0.1% DMSO, tunicamycin (10 µg/mL), and CCCP (20 µM) for the time points indicated in the study. Tunicamycin and CCCP are dissolved in DMSO. For stress washouts cells were washed once with YPD and after centrifugation (2,000 g, 2 min) were resuspended in YPD. For depletion of Not1-AID*-6HA and Scp160-AID*-9myc cells were treated with 100 µM (indole-3-acetic acid) for 1 h and 2 h, respectively. The IAA treatment was kept during stress applications and stress washouts. For dissolution of PBs, cells were treated for 10 min with 1 M 1,6-hexandiol after stress application, the stress was kept during that treatment. To inhibit translation using cycloheximide (CHX), cultures that were treated for 10 min with 100 µg/ml CHX.

### Growth curves and drop tests

Yeast cells were cultured and stressed as described above. For drop tests the culture was diluted to OD_600_ 0.1 and a three 1:10 serial dilutions were made. Then single drops of each condition and dilution were transferred on to a YPD plate and incubated for 48 h at 30°C. Images were acquired using white light mode of the Vilber E-box. For growth curves the stress was applied as described in the according experiment and OD_600_ was measured after 10 min, 2 h, 4 h, 6 h, 8 h, and 21 h.

### Fluorescence microscopy

For live-cell imaging yeast were cultured as mentioned above and resuspended in HC-medium keeping the stress and/or drug treatments present. Images were acquired using Hamamatsu C13440 ORCA.flash4.0 camera mounted on Axio Imager.M2 fluorescence widefield microscope and a X-Cite XYLIS illumination system from excelitas technologies.

PBs were counted manually, when comparing different stress conditions / mutants using ImageJ for Figure 1 and Figure 5. For stress washout experiments as well as experiments using Not1-AID*-6HA, semi-automated approach was used. For that, images were converted to 8-bit in ImageJ, gaussian blur was applied (radius =1). A threshold was set using MaxEntropy, threshold values were minorly adjusted, if needed. The detected PBs were counted and plotted per cell. Fluorescence intensity measurements of PBs were carried out with ImageJ. The mean gray value of the PB was normalized to autofluorescence signal and divided by the normalized mean gray value of a ROI in the cytoplasm. Further, only foci that were 1.2 times brighter than the cytoplasmic background were considered as a PB. Like this, at least 90 PBs from at least three independent experiments were quantified. Colocalization of Dcp2-GFP with other PB components tagged with mCherry were quantified manually for at least 30 cells in three independent experiments.

### Fluorescence recovery after photobleaching

Selected 30-60 min old PBs, marked with Dhh1-GFP, were photobleached for 10 ms using 60% 473 nm laser. Videos were obtained using the Olympus SpinD (CSU-W1) with Photomanipulation/FRAP using an exposure time of 40 ms and 20% 488 nm laser. In total, 451 frames of 0.2 sec per frame were acquired.

Fluorescent intensities of PBs over time were manually analyzed in ImageJ. The fluorescence intensity of the cytoplasm and the intensity of the PB in each frame was measured. The frame of the PB was adjusted as it moves through the cell. During photobleaching of the PB, cytoplasm is bleached as well. To account for this, we measured the intensity of the cytoplasm before and after photobleaching. The recovery intensities were normalized to the average of 15 frames of PB intensity pre-bleaching and the bleaching of the cytoplasm. PBs that were not continuously visible for at least 30 s, were excluded from analysis, unless the PB reappeared at the same position within the following five frames (1 s). For analysis, the data was uploaded to the open source FRAP application FrapBot (http://frapbot.kohze.com). From this we retrieved values for mobile fraction (Mf) and half time of recovery (t_1/2_). Using this approach, we obtained at least 19 PB measurements per condition across three independent experiments.

### L-HPG translation assay

To measure global translation by labeling nascent peptide synthesis, we used the “Click-iT HPG Alexa Fluor 488 Protein Synthesis Assay Kit” (Thermo Fisher Scientific). This assay is based on incorporation of the methionine analog L-HPG that contains an alkyne moiety, to which the fluorescent picolyl azide AF488 can be clicked. The fluorescent intensity is a direct read-out of levels of protein synthesis. For that, cells were cultured in HCD medium (20 µg/µL adenine hemi-sulfate (Sigma A-3159), 35 µg/µL uracil (Sigma U-0750), 80 µg/µL L-tryptophan (Roth 4858.2), 20 µg/µL L-histidine-HCl (Sigma H5659), 80 µg/µL L-leucine (Roth 3984.1), 120 µg/µL L-lysine-HCl (Sigma L-5626), 20 µg/µL L-methionine (Roth 9359.1), 60 µg/µL L-tyrosine (Roth T207.2), 80 µg/µL L-isoleucine (Roth 3922.2), 50 µg/µL L-phenylalanine (Roth 4491.1), 100 µg/µL L-glutamic acid (Sigma G-5889), 200 µg/µL L-threonine (Roth T206.2), 100 µg/µL L-aspartic acid (Sigma A-7219), 150 µg/µL L-valine (Roth 4879.1), 400 µg/µL L-serine (Roth 4682.1), 20 µg/µL L-arginine-HCl (Sigma A-5131), 6.7 g/L Yeast Nitrogen Base w/o amino acids and 2 % glucose) O/N and were inoculated in the morning at OD_600_ 0.1 and grown until OD_600_ 0.6 -0.8. Then, 1 x 10^7^ cells were spun down (2,000 g, 2 min.) and incubated in HC-Met medium and 50 µM L-HPG at 30°C for 10 min (shaking) with the stressor present. This was followed by a 1x PBS wash and fixation in 70 % ice-cold ethanol for 30 min on ice. The cells were washed with 1x PBS and resuspended in 1% Triton-X100 in PBS and permeabilized at RT on an end-over-end rotator for 20 min. After washing with 3% BSA in PBS cells were resuspended in the Click-iT reaction cocktail (2 mM CuSO_4_, 20 µM AF488 picolyl azide, 1:10 reaction buffer additive in PBS) and incubated for 30 min in the dark. Subsequently, cells were washed 4x with 3% BSA in PBS and 2x with 1x PBS. Cells were resuspended in PBS and measured by FACS. The median fluorescence intensity was computed over 20,0000 events and was normalized to the median fluorescence intensity of untreated cells.

### Total protein extracts and western blot analysis

To look at eIF2*𝛼 phosphorylation by western blot, we used* total protein extracts. Equivalents of 5 units of OD_600_ of cells were spun down, washed and flash frozen and resuspended in 200 µL 1x Laemmli buffer with 1x Halt Protease and Phosphatase inhibitor (Thermo Fisher Scientific). The cells were lysed by adding 100 µL acid-washed glass beads (0.25-0.5 mm) and bead-beating (4°C, 40 s, 6.5 m/s), followed by denaturation for 10 min at 65°C. Cellular debris was pelleted and 13 µL of the supernatant was loaded on a 7-18 % SDS-PAGE. The proteins were blotted onto Amersham Protran Premium 0.45 µM nitrocelluose membrane. Membranes were blocked for 1 h with TBST (20 mM Tris, 150 mM NaCl, pH 7.6 and 0.1% Tween20) with 5% non-fat milk. The primary antibodies were incubated at 4°C overnight. The following antibodies were used: rabbit anti-eIF2*𝛼* total (1:1,000 in 5% non-fat milk, kind gift from Tom Dever), rabbit anti-phospo-eIF2*𝛼 (Ser51) (1:1,000 in Can Get Signal^TM^ Solution 1 (Toyoba), Cell Signalling technology, #9721) and mouse anti-Pgk1 (1:20,000* in 5% non-fat milk*, 22C5D8, Invitrogen, #459250, Thermo Fisher Scientific). After washing, the membranes were incubated with HRP-conjugated secondary antibody (1:10,000 anti-rabbit, Sigma Aldrich A054) in TBST or Can Get Signal^TM^ Solution 2 for 2 h. Again, after washing, the blots were developed using* SuperSignal® West Pico Chemiluminescent Substrate (Pierce). Images were acquired on a FusionFX system (Vilber Lourmat).

Alternatively, to check for auxin-induced depletion, we extracted proteins from 5 units of OD_600_ cells using trichloroacetic acid (TCA) precipitation. The SDS-PAGE, transfer and blocking was carried out as described above. We used the following antibodies to check for Not1-AID*-6HA and Scp160-AID*-9myc degradation: mouse anti-HA (1:1,000 in 5% non-fat milk, 12CA5, Roche, Cat. No. 11 583 816 001) and mouse anti-myc (1:2,500 in 5% non-fat milk, 9E10, Sigma Aldrich M4439). *After washing, the membranes were incubated with HRP-conjugated secondary antibody (1:10,000 anti-mouse, Invitrogen 31430) in TBST. Detection was performed as described above*.

### Cell growth and lysis for Hac1 splicing assay

The cells were grown and treated with the stressors as described above. The positive control is tunicamycin which is used at a concentration of 1 μg/ml for 60 min as described in (*47*). The cells were lysed with 500 μl RIPA buffer in a 1.5 ml microcentrifuge tube (50 mM Tris-HCl pH 8, 150 mM NaCl, 1% v/v NP40, 0.5% w/v sodium deoxycholate, 0.1% w/v SDS, 1 mM PMSF, 2 mM DTT, RNasein 40 U/ml) in a multishaker with 0.5 mm diameter beads (Biospec products-Cat. No. 11079105) for 15 min. The lysate was spun at 2,000 g for 2 min. The supernatant was collected in fresh tubes and followed by two rounds of centrifugation, at 8,000 g first for 5 min and then for 10 min. This supernatant was subject to RNA isolation by hot-phenol method.

### RNA isolation

The supernatant was diluted to 400 μl by adding nuclease free water and 0.3 M sodium acetate. 400 µL phenol (a mixture of 25:24:1 phenol: chloroform: isoamylalcohol) was pre-heated at 65°C for 5 min. The supernatant was added to the pre-heated phenol mix and incubated at 65°C for 5 min with intermittent vortexing each minute and then put on ice for 5 min. Then 2 µL glycogen was added followed by 80 μl chloroform, vortexed to disrupt the phase separation and the samples were centrifuged (10,000 g, 10 min, RT). The aqueous phase was collected in a fresh tube, equal volume of phenol was added, vortexed and centrifuged under same condition. Again, the aqueous phase was collected, this time equal volume of chloroform was added, vortexed and centrifuged under same condition. Finally, the aqueous phase was taken, 2 µL glycogen was added, 1:10 sodium acetate, 2.5 volumes of ethanol, vortexed and stored at -80°C overnight. The next day, the tubes were centrifuged at 16,000 g, 4°C, 30 min, supernatant discarded, the pellet further washed with 70% ethanol, again centrifuged under same conditions for 10 min. This supernatant was removed, and the pellet was left to air dry, followed by resuspension in 25 µL nuclease free water. The RNA concentrations were measured by Nanodrop.

### cDNA synthesis from Hac1 mRNA

In PCR tubes, 1.5 µg of RNA, 1 µL of oligo dT (stock concentration is 500 ng/µL from Promega, REF-C1101), and 1 µL of dNTP mix (stock concentration 10 mM) were added followed by RNase-free water to bring volume to 13 µL. The mix was incubated (65°C, 5 min) and subsequently placed on ice. After a quick spin, 4 µL 5’ First-strand buffer and 2 µL DTT (stock concentration is 0.1 M) was added. After gentle mixing, the samples were incubated (42°C, 2 min). Then 1 µL of SuperScript II reverse transcriptase was added and then incubated (42°C, 50 min) for reverse transcription. Reverse transcription was inactivated (70°C, 15 min). Finally, 180 µL of 10 mM Tris–HCl, pH 8.0 was added to the mix. ThermoFisher SuperScript II reverse transcriptase kit was used with Catalog number: 18064014 (It includes SuperScript II reverse transcriptase, 5’ first-strand buffer and 100 mM DTT)

### Checking HAC1 mRNA splicing using PCR

For PCR amplification, NEB Phusion high-fidelity DNA polymerase kit was used (Cat. no. E0553) A 50 µL PCR mixture was setup by mixing 10 µL of 5X Phusion HF Buffer, 1.5 µL of DMSO, 1 µL of 10 mM dNTP mix, 1 µL each of the forward and reverse primers at stock concentration (10 µM), 5 µL of diluted cDNA, 30 µL of water and 0.5 µL of Phusion High-Fidelity DNA polymerase. The primers are listed in Table S2. The PCR samples were examined by electrophoresis on a 1% agarose gel. The gel was run at 120 V for 40 min. The unspliced HAC1 mRNA is 969 bp while the spliced HAC1 mRNA is 717 bp.

### Coating of 384-Well Plates with Concanavalin A (ConA)

Twenty microliters of a 1 mg/ml ConA solution in water were pipetted into each well of a 384-well plate. After a short 1-min incubation, the ConA solution was completely removed using a vacuum pump. The plate was then left to dry for at least 1 h and up to 24 h before imaging.

### Automated Imaging of Yeast Cells and AI-assisted Quantification of PBs

Yeast cells were grown overnight in YPD at 30°C. Cultures were diluted to an OD_600_ of 0.1 and grown to OD_600_ 0.6-0.8. Then, 100 µl were pelleted and resuspended in 1 ml of the respective HC medium in the presence of the stressor. Cells were mixed by vortexing, and 20 µl of the culture were pipetted into a 384-well plate. Cells were pelleted in the plate for 30 s at 50 rcf.

Imaging of the mCherry and/or GFP signal was performed on an inverted Nikon Ti2 widefield fluorescence microscope using a 100× oil objective (Nikon CFI Plan Apochromat DM Lambda 100x Oil, NA = 1.45, W.D. 0.13 mm) and an ORCA-Fusion digital CMOS camera (Hamamatsu, C14440-20UP), controlled by NIS software. An automated imaging pipeline (Nikon JOBS) was used to sequentially image 12 wells, acquiring 4 images per well, and repeating the sequence every 10 min for 150 min. The first image was acquired 10 min after resuspending the cells in the respective stress condition. During sample preparation, the timing required to image one well was accounted for to ensure that all wells were imaged at the same timepoint after treatment. Images were analyzed using Nikon NIS GA3 software and its AI modules (segment.ai, segmentobjects.ai, denoise.ai). Images were first denoised with the denoise.ai module. A subset of images from different conditions was then used to train two AI modules: one for identifying PBs and one for cell segmentation using the background GFP signal. These modules were applied to automatically segment cells and PBs in all images and to quantify the number of PBs per cell. To remove falsely identified structures, segmented objects were filtered based on their intensity relative to the mean background intensity of their respective cell (e.g., only P bodies at least 20% brighter than the mean background were counted). The appropriate threshold was determined manually for each strain and kept constant across all timepoints within the same imaging session. The following thresholds were used: Dcp1-mCherry/Dcp2-GFP: 5% for mCherry & 25% for GFP; Edc3-mCherry/Dcp2-GFP: 15% for mCherry and 25% for GFP; Dhh1-mCherry/Dcp2-GFP: 20% for mCherry & 25 % for GFP; Xrn1-mCherry/Dcp2-GFP: 5 % for mCherry and 25% for GFP; Pat1-mCherry/Dcp2-GFP: 10% for mCherry & 25% for GFP; Lsm4-mCherry/Dcp2-GFP: 10% for mCherry & 25% for GFP; Dcp2-GFP alone: 30-40%; Edc3-GFP alone: 40-45%; Dcp1-GFP alone: 20-35%.

### Protein expression and purification (Dhh1)

Recombinant Dhh1 was purified as previously described for other DEAD-box ATPases (*48*). Briefly, Dhh1 was expressed in *E. coli* Lemo21 (DE3). Cultures were grown at 37 °C in Terrific Broth (TB) to an OD₆₀₀ of 0.6 and induced with 200 µM IPTG. Expression proceeded overnight at 18°C, and cells were harvested the next morning. Pellets were resuspended in lysis buffer (50 mM Tris-HCl pH 7.5, 1 M NaCl, 10% (w/v) glycerol, protease inhibitors, DNase (Roche, Cat# 10104159001), and RNase (Macherey-Nagel, Cat# 74050)) and lysed by pressure homogenisation using an EmulsiFlex (Avestin), followed by sonication. Initial capture was performed by immobilized metal affinity chromatography (IMAC), followed by overnight 3C protease cleavage and a subsequent reverse IMAC. Final polishing was performed by size exclusion chromatography (SEC) using a 16/600 HiLoad Superdex 200 pg column (Cytiva) on an ÄKTA pure (GE Life Sciences), and the protein was exchanged into storage buffer (1,000 mM NaCl, 50 mM phosphate buffer pH 7.5, 2 mM MgCl₂, 3 mM 2-mercaptoethanol, and 10% glucose). Purified protein was snap-frozen in liquid nitrogen and stored at -80 °C.

### Chemical labeling of proteins

Untagged Dhh1 was incubated for at least 1 h in the dark with ATTO565 NHS-ester dye (ATTO-TEC GmbH, Cat# AD 565-31) at a five-fold molar excess over the protein. To preferentially label N-terminal amines, the reaction pH was adjusted to 6.5. Free dye was removed using Zeba Spin Desalting Columns (Thermo Fisher), and labelled protein was concentrated to 100-200 µM using Amicon Ultra-4 centrifugal filters (Merck Millipore).

### *In vitro* condensation assay

Dhh1 condensation assays were performed in 384-well, black, optically clear, flat-bottom, ultra-low-attachment-coated PhenoPlates (Revvity, Cat# 6057800) as previously described (*48*). Briefly, 2 µL of 50 µM Dhh1 in storage buffer, spiked with 1% labelled protein, was dispensed to the side of each well and mixed with 18 µL master mix to reach final assay conditions of 50 mM sodium phosphate (pH 6.4), 100 mM NaCl, 2 mM DTT, 0.05 mg/mL BSA, and 2 mM MgCl_2_, supplemented with the indicated RNA concentrations. Poly(U) (Cat# P9528-25 mg) and Poly(A) (Cat# P9403-25 mg) RNA were obtained from Sigma. After reaction assembly, plates were centrifuged briefly (30 s at 100 rcf) and incubated for 45 min at 25 °C prior to imaging. Images were acquired on a temperature-controlled (25 °C) inverted Nikon Ti2 microscope equipped with a Crest X-Light V3 spinning disk module, a 40X Plan Apo Lambda air objective (NA 0.95), a Lumencor Celesta light source, and a Teledyne Photometrics Kinetix cMOS camera. The microscope was controlled using Nikon NIS-Elements software and automated via the integrated JOBS system. Condensates were segmented using CellProfiler (v4.2.6). For each droplet, the partition coefficient (brightness) was calculated as the ratio of the median fluorescence intensity inside the condensate to the median fluorescence intensity of the immediate surrounding.

### Statistical Analysis

All statistical analysis was performed in Graph Pad Prims Version 10.6.1 (799), and p < 0.05 was considered statistically significant. Data was tested for normal distribution. Statistical analysis was assessed either for multiple comparisons by analysis of variance (ANOVA) with post-hoc Dunnett’s, Tukey’s or Sidak’s test, as appropriate, or two-tailed Welch’s t-test for two component comparison analysis. The number of independent biological replicates are indicated in the figure.

## Acknowledgments

We are grateful for Tom Dever for antibodies.

## Funding

Marie Sklodowska-Curie Postdoctoral Fellowship (101030442) to D.M. Swiss Life to D.M.

Grant for junior researcher from the University of Basel to D.M.

Swiss National Science Foundation (310030_185127, 310030_197779, 310030_219513) to A.S

University of Basel to A.S.

## Author contributions

Conceptualization: AS, DM, MR

Methodology: DM, MR, FW, MS, AS

Investigation: DM, MR, FW, MS

Visualization: DM, MR, FW, MS

Supervision: AS, MH

Writing—original draft: DM, MR, AS

Writing—review & editing: DM, MR, FW, MS, AS, MH

## Data and materials availability

All data are included in the manuscript.

---

**Figure S1.**
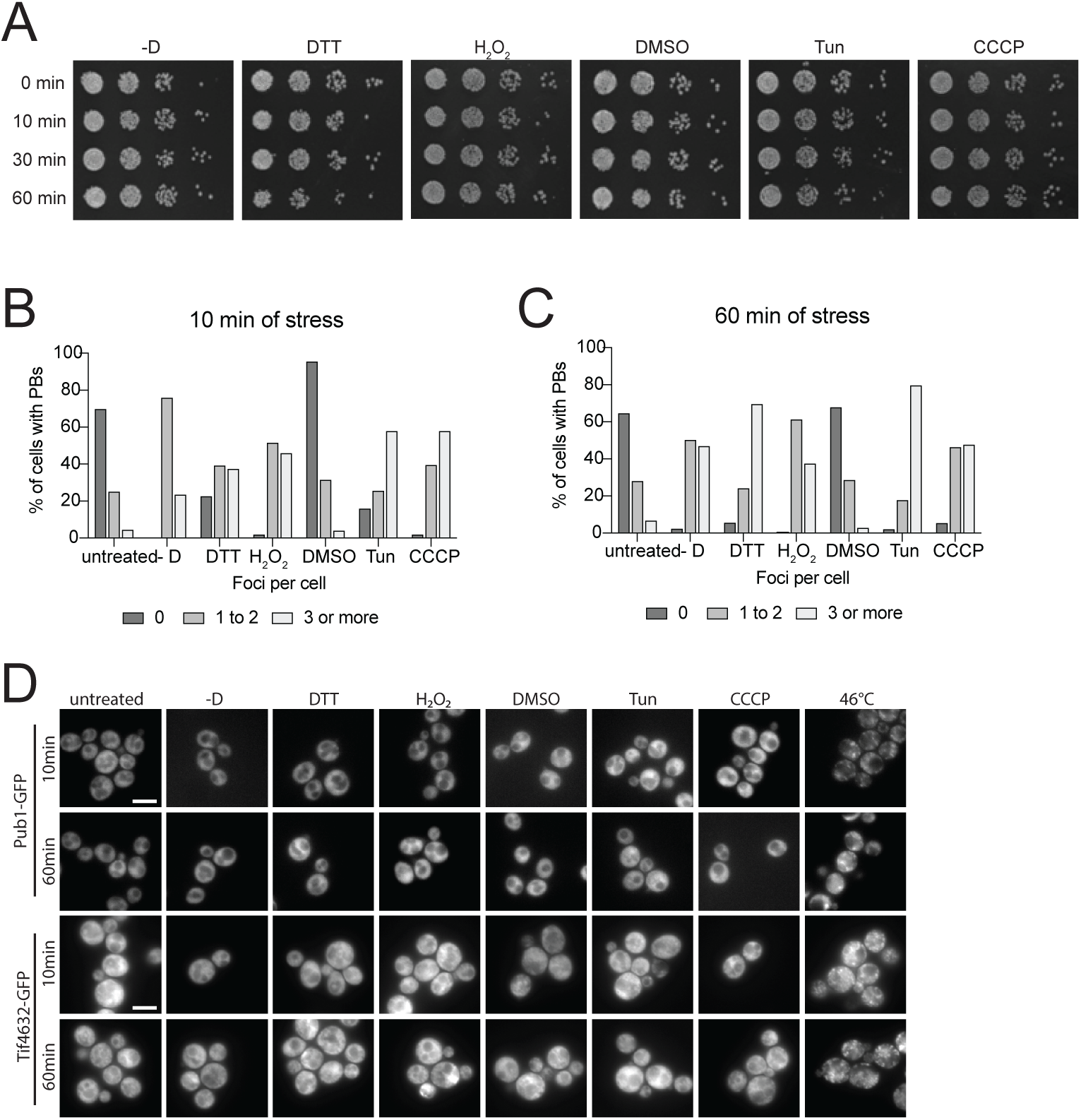
Stress repertoire is not lethal for the cells and does not induce SG formation. (A) Yeast cells were treated with the indicated stresses for the specified time points, spotted in 10-fold serial dilutions on YPD, and grown at 30 °C. n=3. (B) Binning of PBs per cell in the different stress conditions after 10 and (C) 60 min. DTT and tunicamycin show later onset of PB formation in 15-20% of the cells. n=3. (D) Imaging of SG formation (Pub1-GFP and Tif4632-GFP) after 10 and 60 min of stress treatment. n=3. Scale bar = 5 µm.

**Figure S2.**
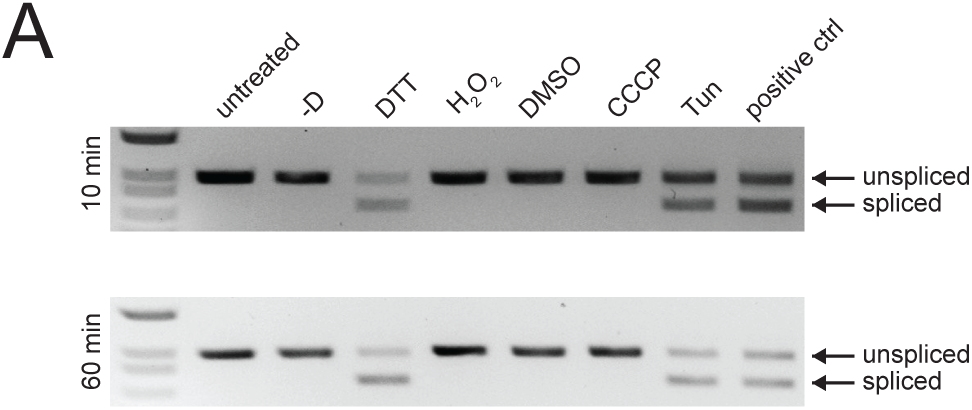
UPR is activated in DTT- and tunicamycin-treated cells. (A) Visualization of spliced (717 bp) and unspliced (969 bp) HAC1 mRNA by semi-quantitative PCR and gel electrophoresis across all stresses from the stress repertoire. n=3.

**Figure S3.**
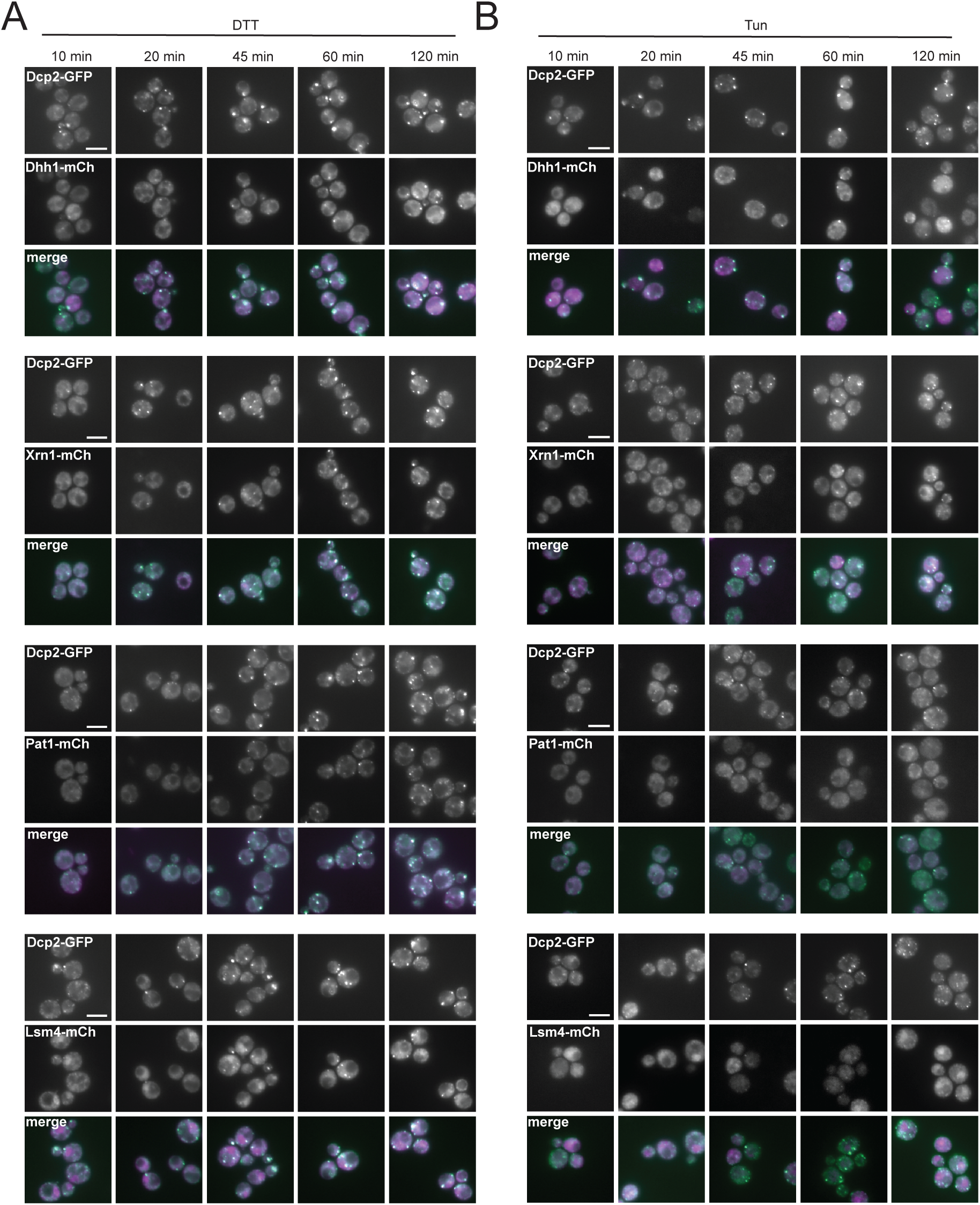
Sequential recruitment of PBs induced by DTT and tunicamycin treatment. (A) Imaging of recruitment of mCherry-tagged Dhh1, Xrn1, Pat1 and Lsm4 to Dcp2-GFP foci upon DTT treatment and (B) tunicamycin treatment. n=3. Scale bar = 5 µm.

**Figure S4.**
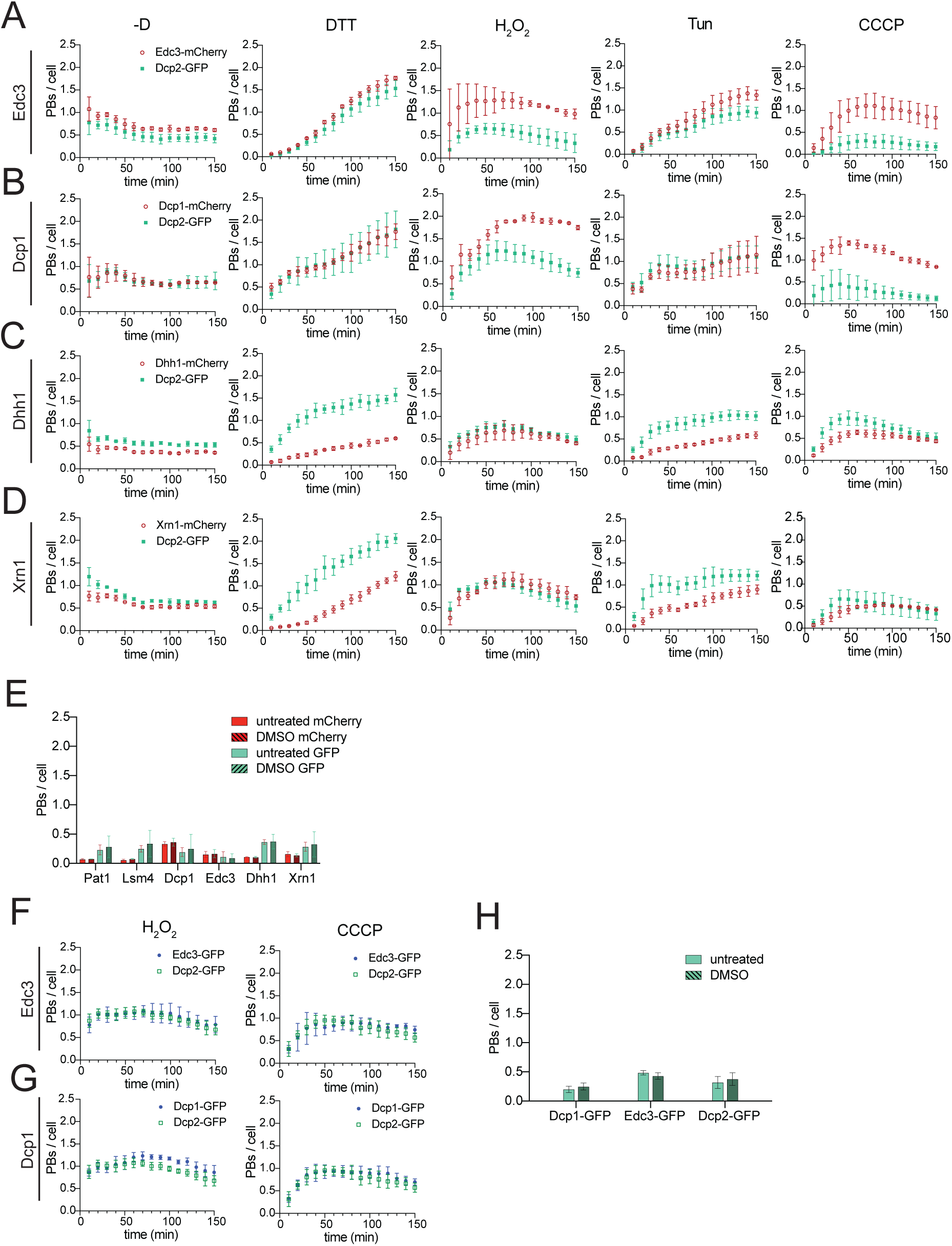
Automated imaging and AI-assisted analysis confirms staggered recruitment of PB at different stresses. (A-D) Q**uantification of PB/cell for double tagged strains expressing Dcp2-GFP together with either Edc3, Dcp1, Dhh1 or Xrn1 tagged with mCherry.** Cells were imaged automatically in 384-well plates at the indicated timepoint and stress, and PBs were quantified using Nikon NIS AI modules. Data represent the mean identified number of PB/cell ± standard deviation from three independent biological replicates, with ≥250 cells quantified per time point and n=3 independent replicates. (E) Quantification of PBs per cell under control conditions (untreated for glucose, H_2_O_2_, and DTT stress; DMSO-treated for CCCP and tunicamycin) in double-tagged strains expressing Dcp2-GFP together with either Edc3-, Dcp1-, Dhh1-, or Xrn1-mCherry. Data represent the mean number of PBs/cell averaged across all time points under the same control condition ± standard deviation from n=3 independent biological replicates. (F,G) Quantified PB/cell of single tagged strains expressing either Edc3-GFP or Dcp1-GFP compared to a strain expressing Dcp2-GFP treated with H_2_O_2_ or CCCP confirming similar recruitment dynamics. Cells were imaged automatically in 384-well plates at the indicated timepoint and stress, and PBs were quantified using Nikon NIS AI modules. Data represent the mean identified number of PB/cell ± standard deviation from n=3 independent biological replicates, with ≥250 cells quantified per time point and replicate. (H) Quantification of PBs per cell under control conditions (untreated H_2_O_2_; DMSO for tunicamycin and CCCP) for strains expressing either Edc3-GFP, Dcp1-GFP or Dcp2-GFP. Data represent the mean number of PBs/cell averaged across all time points under the same control condition ± standard deviation from three independent biological replicates.

**Figure S5.**
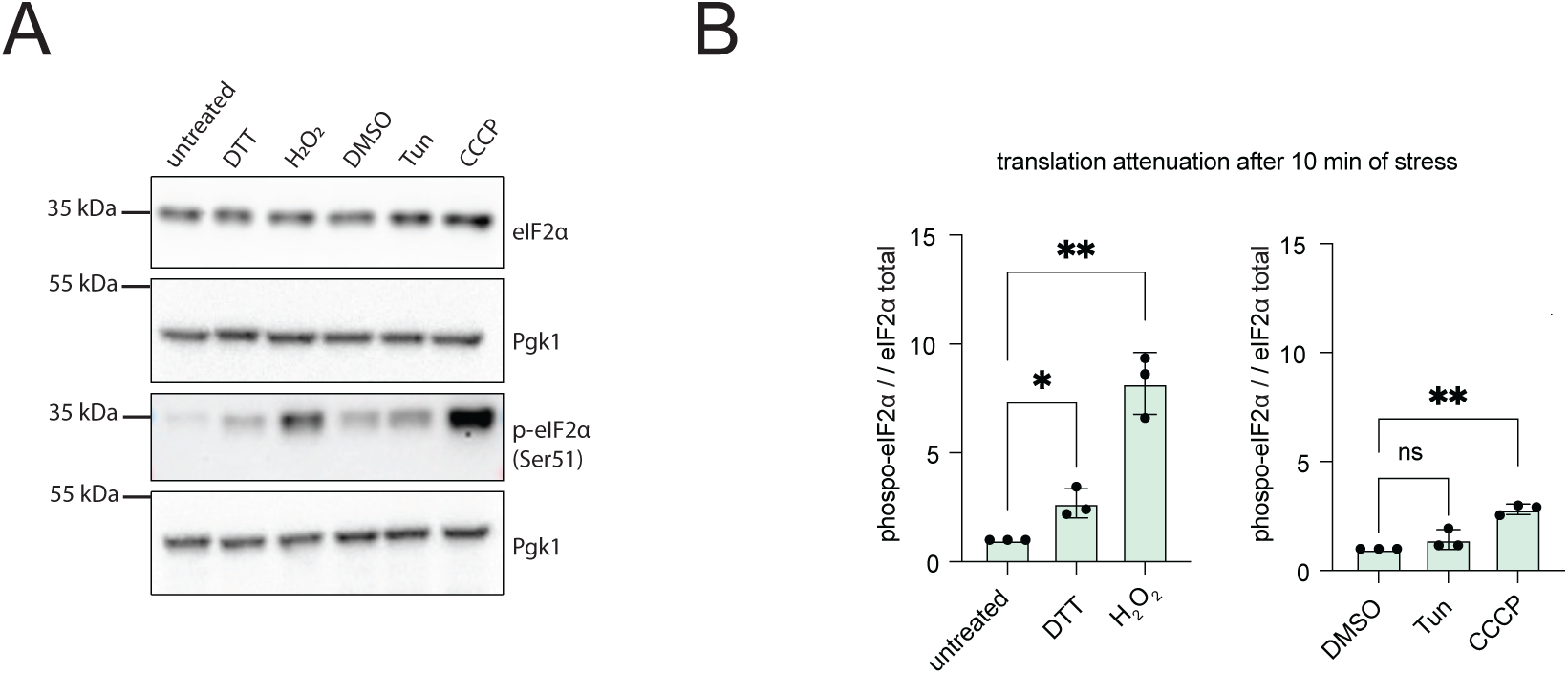
Translation initiation is attenuated after 10 min of DTT, H_2_O_2_ and CCCP, but not tunicamycin treatment. (A) Western blot of total eIF2α and phospho-eIF2α (Ser 51) after 10 min of stress. Pgk1 was used as loading control. n=3. (B) Phosphorylation levels were quantified as the ratio of phosphorylated to eIF2α total protein for each n=3 biological replicate. For visualization, data are normalized to the mean of the untreated / DMSO control. Statistical analysis was performed on the log2-transformed raw phospho/total ratios using one-way ANOVA followed by Dunnett’s multiple comparisons test comparing each treatment to untreated control. Data are presented as mean ± SD (n = 3 biological replicates).

**Figure S6.**
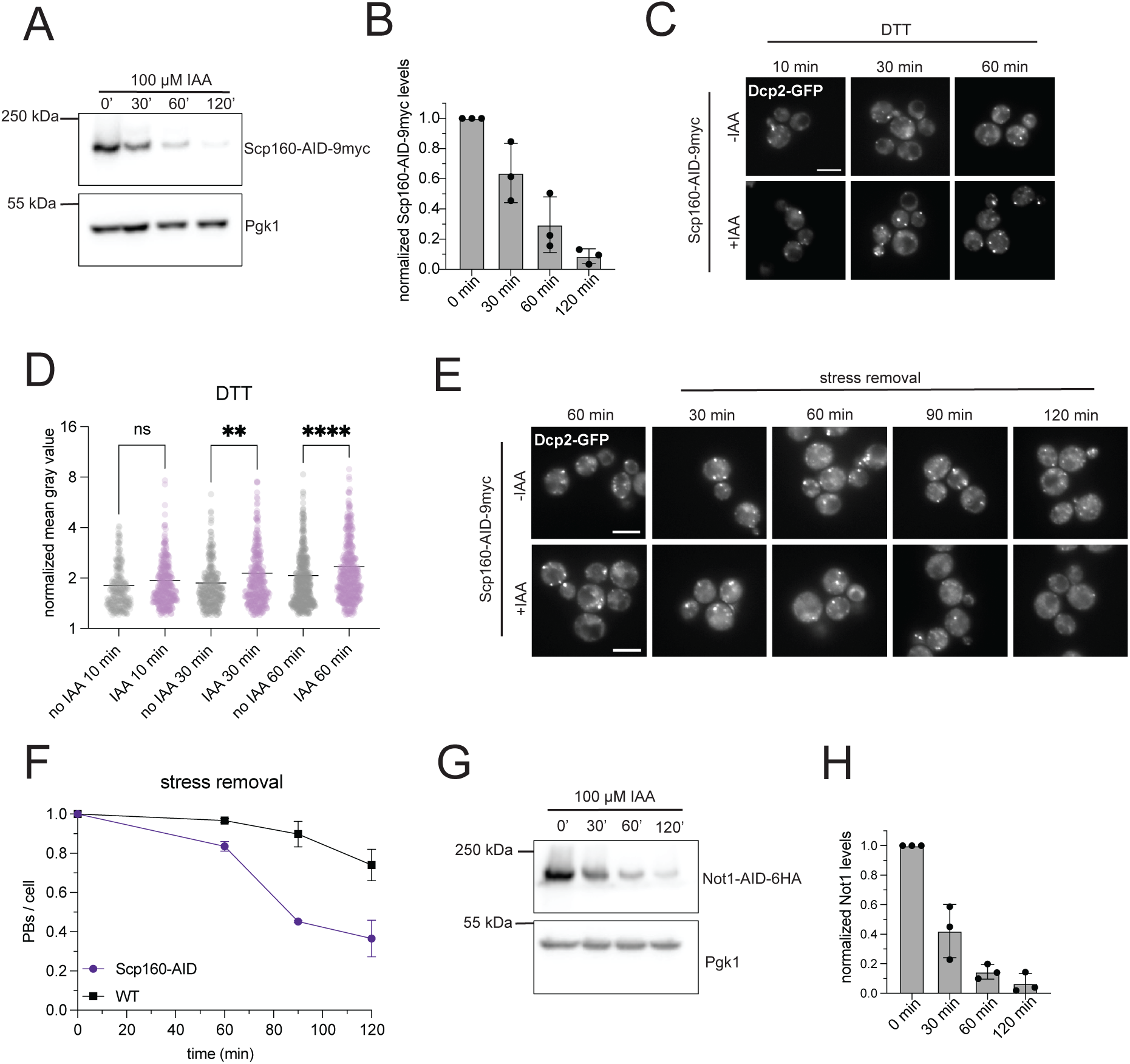
Depletion of Scp160 and Not1 lead to changes in properties of DTT PBs. (A) Western Blot of the depletion of Scp160-AID*-9myc after 120 min of 100 µM IAA treatment. Pgk1 was used as loading control. (B) Quantification of the relative changes of Scp160-AID-9myc levels upon 100 µM IAA shown in (A). n=3. One-way ANOVA with Tukey’s multiple-comparisons test was used for statistical analysis. (C) Imaging of DTT induced PBs (Dcp2-GFP) in Scp160-depleted (100 µm IAA) and non-depleted cells after 10, 30 and 60 min of stress induction. n=3. Scale bar = 5 µm. (D) Quantification of the fluorescence intensity of DTT PBs captured in (C). The y-axis is plotted in log_2_ scale. Statistical analysis was performed using a Kruskal-Wallis test with Dunn’s multiple-comparisons test. (E) Kinetics of PB (Dcp2-GFP) dissolution after DTT stress removal in Scp160-depleted (+IAA) and control cells (-IAA) at indicated timepoints. n=3. Scale bar = 5 µm. (F) Quantification of PB number per cell expressed as change relative to 10 min of DTT treatment in Scp160-depleted (+IAA) and non-depleted (-IAA) cells. (G) Western Blot depicting the depletion of Not1-AID*-6HA after 30, 60 and 120 min of 100 µM IAA treatment. Pgk1 was used as loading control. n=3. (H) Relative changes of Not1-AID*-6HA protein levels upon 100 µM IAA shown in (G). n=3.

**Figure S7.**
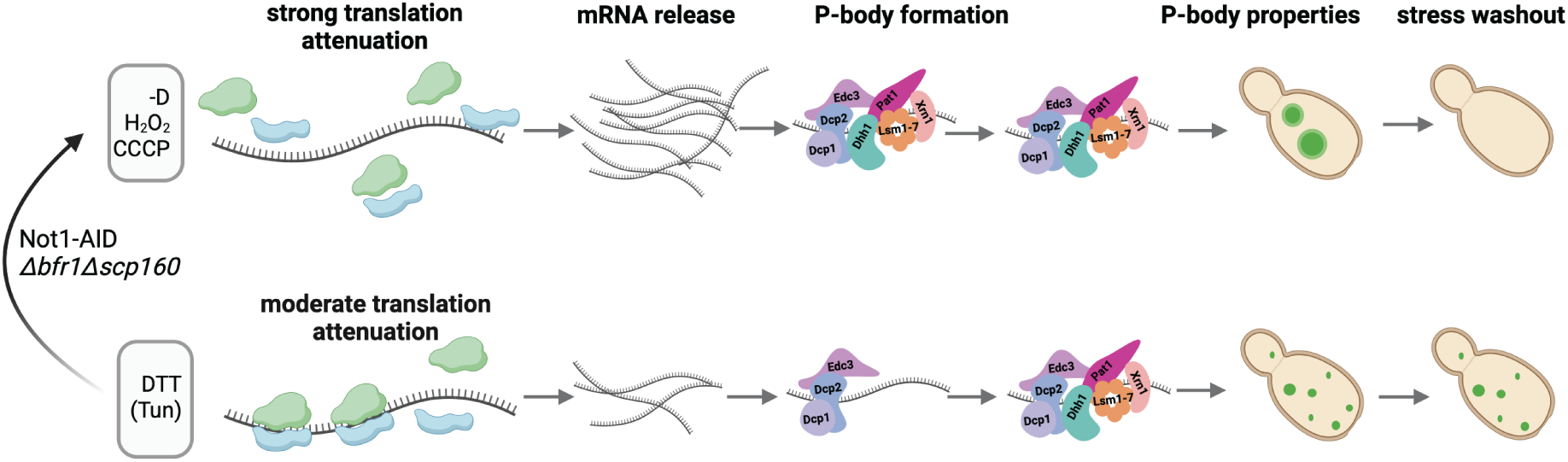
Non-translating mRNA determines PB properties. Depicts the current working model. The amount of translation attenuation correlates with the non-translated mRNA that can be bound by PB core components – the decapping machinery – and assembles into PBs. The brightness, assembly, composition as well as dissolution upon stress washout of these PBs is dependent on the amount of non-translated mRNA. The pool of non-translated mRNAs can be increased by genetic perturbations, namely, deletion of the polysome associated proteins Bfr1 and Scp160 as well as by depletion of the Ccr4-Not deadenylase scaffold Not1 that can then convert persistent PBs to a more temporary PB phenotype.

**Table S1.**
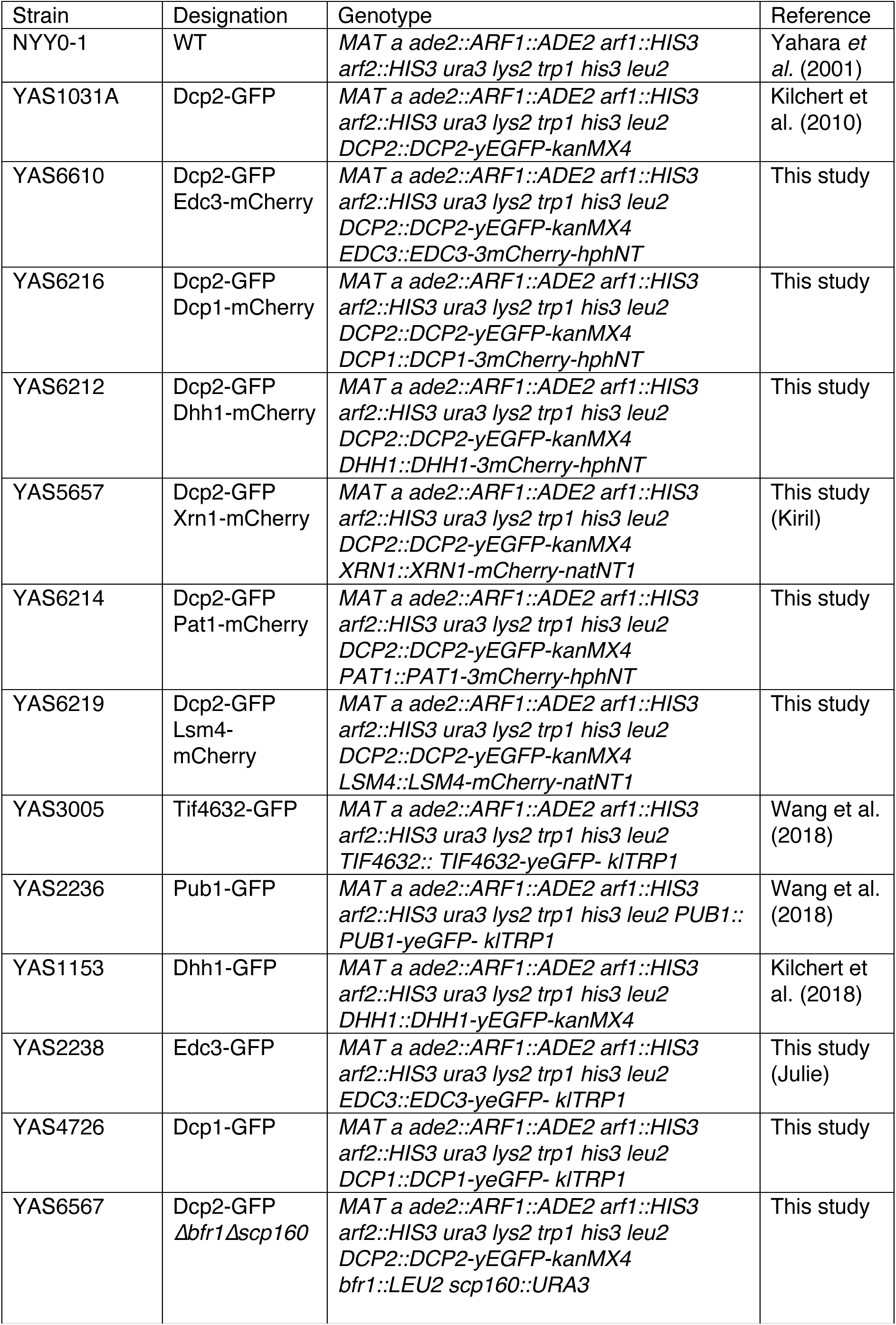
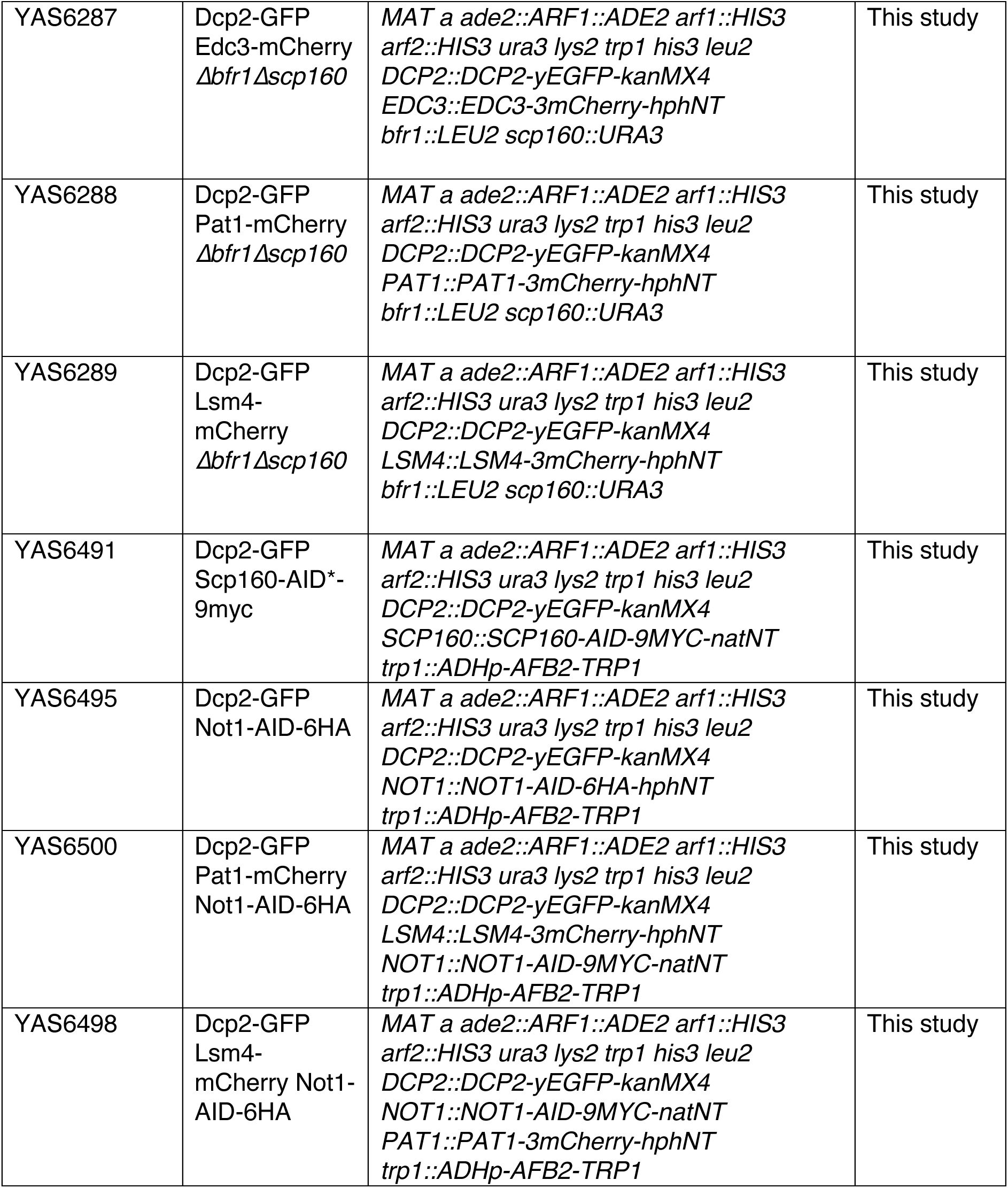
Yeast Strains.

**Table S2.**
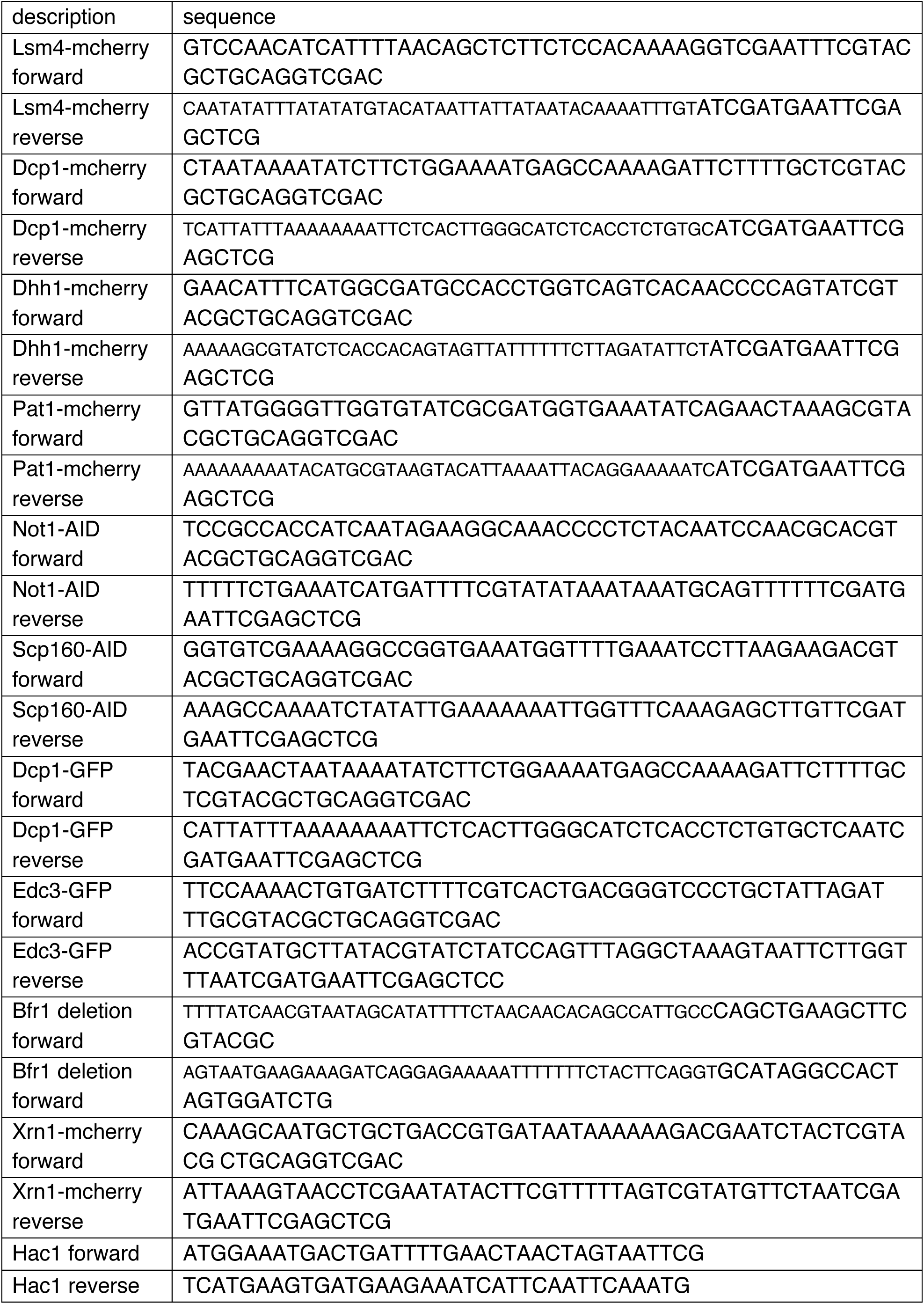
Primer List.

